# Mono-ubiquitination of Rabphilin 3A by UBE3A serves a non-degradative function

**DOI:** 10.1101/2020.12.14.422704

**Authors:** Rossella Avagliano Trezza, A. Mattijs Punt, Edwin Mientjes, Marlene van den Berg, F. Isabella Zampeta, Ilona J. de Graaf, Yana van der Weegen, Jeroen A. A. Demmers, Ype Elgersma, Ben Distel

## Abstract

Angelman syndrome (AS) is a severe neurodevelopmental disorder caused by brain-specific loss of UBE3A, an E3 ubiquitin protein ligase. A substantial number of possible ubiquitination targets of UBE3A have been identified, although evidence of being direct UBE3A substrates is often lacking. Here we identified the synaptic protein Rabphilin-3a (RPH3A), an effector of the RAB3A small GTPase involved in axonal vesicle priming and docking, as a ubiquitination target of UBE3A. We found that the UBE3A and RAB3A binding sites on RPH3A partially overlap, and that RAB3A binding to RPH3A interferes with UBE3A binding. We confirmed previous observations that RPH3A levels are critically dependent on RAB3A binding but, rather surprisingly, we found that the reduced RPH3A levels in the absence of RAB3A are not mediated by UBE3A. Indeed, while we found that RPH3A is ubiquitinated in a UBE3A-dependent manner in mouse brain, UBE3A mono-ubiquitinates RPH3A and does not facilitate RPH3A degradation. Moreover, we found that an AS-linked UBE3A missense mutation in the UBE3A region that interacts with RPH3A, abrogates the interaction with RPH3A. In conclusion, our results identify RPH3A as a novel target of UBE3A and suggest that UBE3A-dependent ubiquitination of RPH3A serves a non-degradative function.

## Introduction

AS is a severe neurodevelopmental disorder characterized by intellectual disability, impaired motor coordination, epilepsy and behavioral abnormalities ^1^. AS is associated with functional loss of the HECT (<u>H</u>omologous to the <u>E</u>6-AP <u>C</u>arboxyl <u>T</u>erminus) E3 ligase UBE3A (also known as E6AP) encoded by the *UBE3A* gene ^1,2^. E3 ligases are responsible for protein ubiquitination, a cellular code that dictates the destiny of most of our proteins ^3^. The fate of the ubiquitinated protein is dependent on the number of ubiquitin (Ub) molecules conjugated and the type of Ub-Ub linkage in the poly-ubiquitin chain. Conjugation of a single Ub molecule to a lysine side chain of the target protein results in mono-ubiquitination, a modification that can change the localization or control the activity of the target. The target-attached Ub can be further modified on its N-terminus or on one of the seven lysine residues, leading to the synthesis of poly-ubiquitin chains with different linkage types. While the function of most of the Ub linkage types have not been firmly established, K48-linked Ub chains, the predominant linkage type in cells, target proteins to the proteasome for degradation (reviewed ^4,5^). Although it has been well documented that UBE3A mainly synthesizes K48-linked poly-ubiquitin chains, recent evidence suggests that the UBE3A HECT domain can synthesize ubiquitin chains with different linkage type specificity ^6^.

In order to understand the molecular basis of AS, several studies have been conducted to identify direct and indirect targets/interactors of UBE3A, but for most of these identified proteins their relevance for the disorder remains unknown ^7^. Furthermore, many of the discovered UBE3A-interacting proteins do not show UBE3A-dependent proteasomal degradation, suggesting that these interacting proteins are either not *bona fide* UBE3A ubiquitination substrates or they receive a type of ubiquitination which does not result in their degradation.

The synaptic vesicle release cycle at the presynaptic bouton forms the basis for signal transmission between neurons ^8^, and alterations in the vesicle cycle often lead to neurological disorders^9^. Previous studies showed an accumulation of clathrin-coated vesicles in a *Ube3a* mouse model of AS ^10^, indicating a role for UBE3A in presynaptic function. However, the mechanistic underpinnings of this impaired endocytosis and its relevance to the disorder is unknown.

In this study we identify the presynaptic protein Rabphilin-3A (RPH3A), as a novel binding partner of UBE3A. RPH3A is an effector protein that binds to the GTP (guanosine triphosphate)-bound form of RAB3A holding the protein in this state by preventing its GTPase activity from hydrolyzing GTP ^11^. Members of the RAB3 family, which are small GTP-binding proteins that cycle between a GTP and GDP (guanosine diphosphate)-bound state^12^, are key regulators of vesicle exocytosis. At least four isoforms of RAB3 have been identified: RAB3A, RAB3B, RAB3C and RAB3D. RAB3A is the most abundant isoform in the brain ^8,12-14^ and the better-described member of the family. RAB3A has been shown to interact with members of the SNARE (<u>SNA</u>P (Soluble NSF[N-ethylmaleimide-sensitive factor] Attachment Protein <u>RE</u>ceptor)) complex to facilitate vesicle fusion ^15^. RAB3A associates with vesicles in its GTP-bound form and is released once the GTP has been hydrolyzed to GDP ^15^. The balance between the GTP- and the GDP-bound form of RAB3A is maintained by the RAB3A GDP-dissociation inhibitor protein (GDI), which keeps RAB3A in the GDP-bound state and promotes its dissociation from synaptic vesicles ^16^.

Detailed analysis of the RPH3A-RAB3A interaction has led to the identification of two regions in RPH3A that are involved in RAB3A binding: i) specific residues in the first α-helix (α-helix a1) of RPH3A (amino-acids 45-161) and ii) the SGAWFF sequence, a motif localized C-terminal of the Zn^2+^ binding domain of RPH3A that is highly conserved among other RAB effector proteins (e.g. RIM, Noc2) ^11^. RPH3A also contains two C2 domains homologous to the C2A and C2B of Synaptotagmin1 (Syt1), which are responsible for Ca^2+^ binding ^17^. While *Rab3a* knock-out (KO) mice have revealed discrete but important functions of this small GTPase in vesicle exocytosis ^14^, deletion of its effector RPH3A failed to show a major impairment^18^. Interestingly, mice lacking RAB3A show a marked decrease in RPH3A levels, suggesting that in the absence of RAB3A the stability of RPH3A is severely affected^14^.

Here we show that UBE3A binds to a N-terminal alpha helical segment of RPH3A and provide evidence that this binding site partially overlaps with the RAB3A binding site, the binding of which is required for RPH3A stability. We further show that RPH3A ubiquitination in mouse brain is strongly dependent on the presence of UBE3A and that RPH3A is a direct ubiquitination substrate of UBE3A in a bacterial ubiquitination assay. UBE3A-dependent ubiquitination of RPH3A does not result in its degradation, and RPH3A levels are not changed in brains of *Ube3a* mice nor in *Ube3a/Rab3a double* mutants. Taken together, these results suggest that the ubiquitin modification of RPH3A by UBE3A serves a proteasome-independent function.

## Results

### Identification of RPH3A as a binding partner of UBE3A

In a previous study we screened for proteins that interact with UBE3A using a yeast two-hybrid (Y2H) screen ^19^. One of the four high confidence interacting proteins found in this screen was Rabphilin3A (RPH3A). RPH3A is a RAB3 effector ^11^ that contains an N-terminal zinc-finger domain and two C-terminal Ca^2+^-binding C2 domains (**Fig. 1**). The clone identified suggested that the first 157 amino acids of RPH3A, which includes the Zn-finger domain, are required for interaction with UBE3A. In order to confirm these results and further delineate the UBE3A binding region, we generated several deletion mutants of RPH3A and tested them for interaction with wild type UBE3A in an Y2H assay. As shown in **Figure 1** (see also **Supplementary Fig. S1**) we were able to confirm that both full length RPH3A and the clone originally found in the screen (RPH3A^1-157^) interact with UBE3A. Interestingly, RPH3A^1-157^ harbors the two domains essential for the interaction with the small GTPase RAB3A ^13,17^: the N-terminal α-helix a1 (amino acid 48-85, secondary structure prediction at PSIPRED: http://bioinf.cs.ucl.ac.uk/psipred/ psipred prediction) and the SGAWFF motif, a strictly conserved motif among GTPase effectors, that is located C-terminally of the zinc finger region in RPH3A ^11^ (**Supplementary Fig. S2**). Consistent with this, RPH3A^1-157^ interacted strongly with RAB3A^Q81L^, a dominant active form of RAB3A that is locked into its GTP bound form ^8,20^ (**Fig. 1)**.

**Fig 1.**
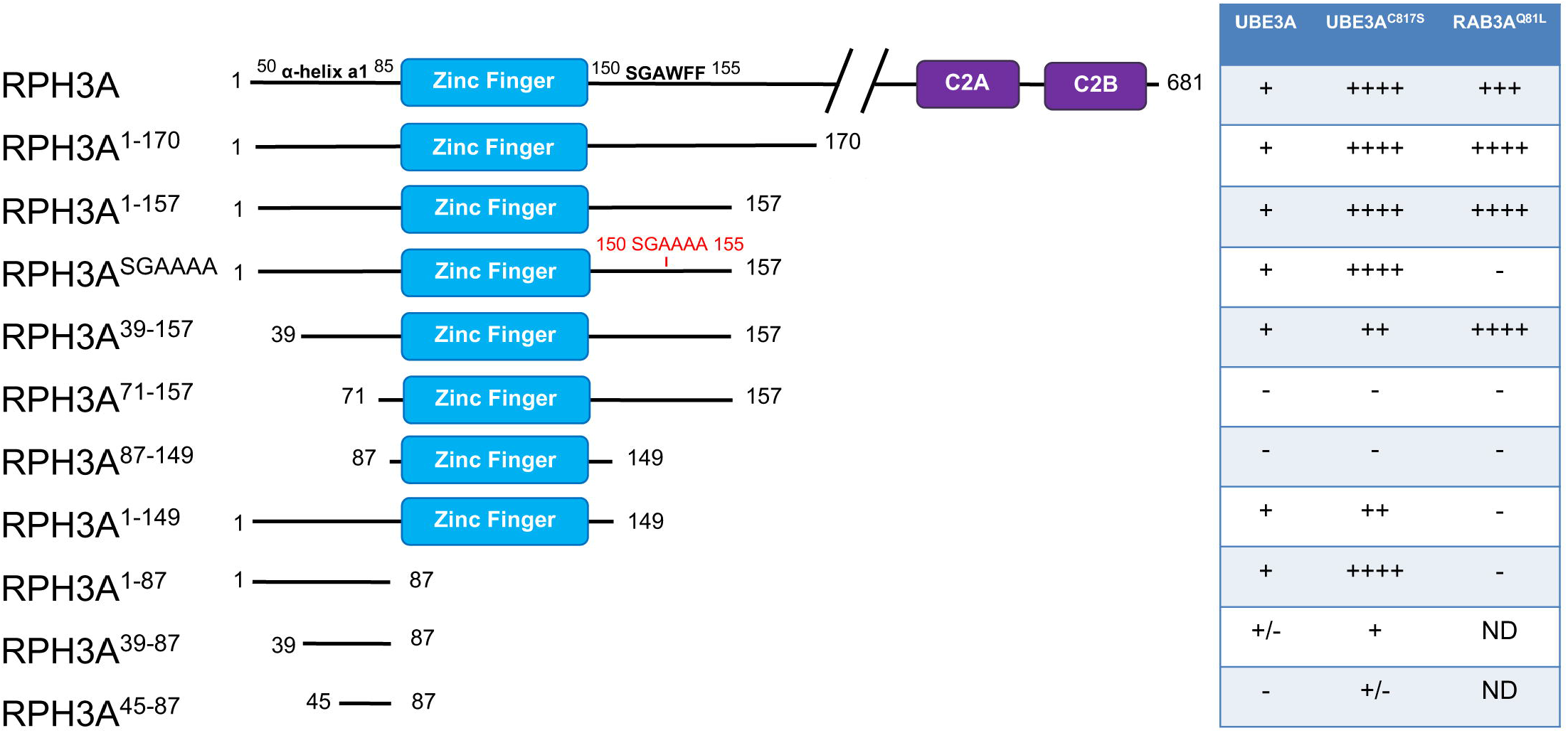
Identification of RPH3A as a novel binding partner of UBE3A. Schematic representation of RPH3A fragments employed in this study and their strength of interaction with wild type UBE3A, catalytically inactive UBE3A (UBE3A^C817S^) and the GTP-locked mutant of RAB3A (RAB3A^Q81L^). *S. cerevisiae* Y2H Gold and Y187 strains transformed with UBE3A, UBE3A^C817S^ or RAB3A^Q81L^ (baits) and RPH3A constructs (preys), respectively, were mated and analyzed for strength of interaction by measuring the activation of the *HIS3* and *ADE2* reporter genes. (-) no interaction (no growth on any of the selection plates); (+) weak interaction (growth His^−^ plates); (++) medium interaction (growth on His^−^ and His^−^ plus 10mM 3-AT); (+++) medium-strong interaction (growth on His^−^ and His^−^ plus 10 and 20mM 3-AT); (++++) strong interaction (growth on all selection plates including Ade^−^). Scans of the selection plates are presented in **Supplementary Figure S1**. Indicated is the Zn finger domain (cyan), the C2 domains (purple) and the SGAWFF motif required for RAB3A interaction. ND = Not determined. Note: RPH3A^39-87^ and RPH3A^45-87^ were only tested on His^+^ and His^-^ selection plates.

To determine whether UBE3A and RAB3A share the same binding sites on RPH3A we first mutated the aromatic residues in the SGAWFF motif of RPH3A^1-157^ to alanine, a mutation that has been previously described to disrupt RPH3A-RAB3A interaction ^17,21^. As shown in **Figure 1**, RPH3A^SGAAAA^ was unable to associate with RAB3A^Q81L^, while its interaction with UBE3A was maintained, indicating that the conserved SGAWFF motif is not required for UBE3A interaction. Deletion of the N-terminal 39 amino acids (RPH3A^39-157^), leaving the α-helix a1 intact, did not affect either UBE3A or RAB3A^Q81L^ binding. However, upon deletion of the complete α-helix a1 (constructs RPH3A^71-157^ and RPH3A^87-157^), the interaction with both UBE3A and RAB3A^Q81L^ was lost. To test whether the RPH3A N-terminal α-helix is sufficient for UBE3A association, we generated RPH3A fragments harboring the α-helix a1 but lacking the entire zinc finger domain. The Y2H experiment showed that the interaction with UBE3A was completely preserved with RPH3A^1-87^ and only partially retained with RPH3A^39-87^ and RPH3A^45-87^. Notably, all UBE3A-RPH3A interactions appeared to be stronger with the active site mutant of UBE3A (UBE3A^C817S^) compared to that with wild type UBE3A, possibly suggesting increased binding in the absence of ubiquitin transfer. The molecular basis for this increased interaction with UBE3A^C817S^ remains to be investigated. Taken together, these results indicate that RPH3A is a *bona fide* binding partner of UBE3A and that the minimal UBE3A binding domain of RPH3A (residues 39 to 87) includes the complete α-helix a1. Furthermore, these results show that UBE3A and RAB3A^Q81L^ have partially overlapping binding sites on RPH3A, the exact nature of which was further investigated in this study (see below).

To investigate if the UBE3A-RPH3A interaction is direct, we performed co-immunoprecipitation (co-IP) experiments using proteins expressed in bacteria. Specifically, *E. coli* cells co-expressing HA-UBE3A and V5-RPH3A or HA-UBE3A^C817S^ and V5-RPH3A, were lysed and an HA immuno-precipitation (IP) experiment was performed. Cells co-expressing HA-GFP and V5-RPH3A were used as negative control. As shown in **Figure 2a**, both UBE3A and UBE3A^C817S^ were able to pull down RPH3A, while the GFP control did not. Similarly, we investigated the ability of shorter constructs of RPH3A to interact with UBE3A. Lysates from *E. coli* cells co-expressing HA-UBE3A and RPH3A^1-157^, RPH3A^39-157^ or RPH3A^71-157^ were subjected to IP with anti-HA or IgG (control) (**Fig. 2b**). Both RPH3A^1-157^ and RPH3A^39-157^ were pulled down by UBE3A with equal efficiency, while RPH3A^71-157^ showed a strongly reduced interaction with UBE3A. Taken together, these results show that the UBE3A-RPH3A interaction is direct and suggest an important role of the first α-helix in RPH3A for UBE3A binding.

**Fig 2.**
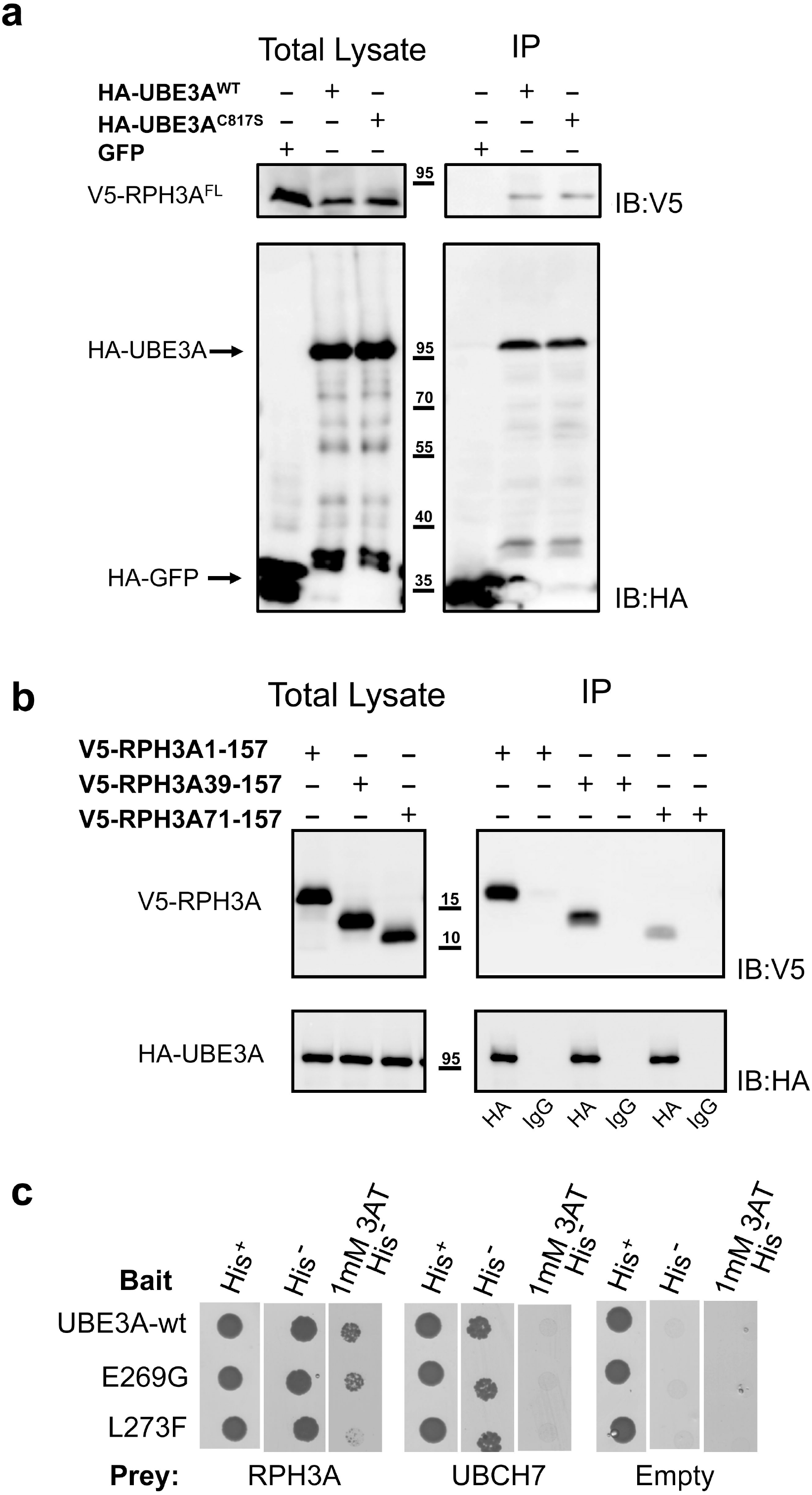
The UBE3A-RPH3A interaction is direct and is disturbed by an AS-linked mutation in UBE3A. **a**) *E. coli* cells co-expressing HA-UBE3A and V5-RPH3A or HA-UBE3A^C817S^ (catalytically inactive UBE3A) and V5-RPH3A, were lysed and cleared lysates were subjected to immunoprecipitation (IP) with anti-HA antibodies. Cells co-expressing HA-GFP and V5-RPH3A were used as negative control. Approximately 4X of the IP fractions, compared to the total lysates, were loaded and analysed by SDS-PAGE and anti-HA and anti-V5 immunoblotting. **b**) *E. coli* cells co-expressing HA-UBE3A and V5-RPH3A^1-157^, V5-RPH3A^39-157^ or V5-RPH3A^71-157^, were lysed and cleared lysates were subjected to IP with anti-HA or IgG (negative control). Approximately 4X of the IP fractions, compared to the total lysates, were loaded and analysed by SDS-PAGE and anti-HA and anti-V5 immunoblotting. Full-length blots are presented in **Supplementary Figure S8. c)** Yeast two-hybrid interaction of RPH3A^1-157^ with wild-type UBE3A and two Angelman syndrome-linked variants (p. E269G and p. L273F). Interaction strength was assessed as described in the legend to Figure 1. Scans of the selection plates are presented in **Supplementary Figure S3a**.

### An AS-linked mutation in the RPH3A binding site of UBE3A interferes with RPH3A binding

In a previous study we showed that the RPH3A binding-region in UBE3A is located in the N-terminus between residues 253 and 274 ^19^. Of note, within this small UBE3A region several AS-linked variants have been identified (https://www.ncbi.nlm.nih.gov/clinvar/?term=UBE3A[gene]). To investigate if AS-linked mutations in this region affect RPH3A binding we introduced two variants, p.Glu269Gly and p.Leu273Phe, in UBE3A by site directed mutagenesis and tested their interaction with full-length RPH3A (**Fig 2c and Supplementary Fig. S3a**). While the strength of interaction of the p.Glu269Gly variant with RPH3A was comparable to wild type UBE3A, the p.Leu273Phe variant showed a reduced interaction with RPH3A. Control experiments showed that the interaction of the p.Leu273Phe variant with UBCH7 (cognate E2 of UBE3A) is undisturbed and that the expression of the mutant protein is comparable to wild type UBE3A (**Fig. 2c and Supplementary Fig. S3a and S3b**).

### Identification of the RPH3A residues involved in UBE3A and RAB3A interaction

Given that the first α-helix in RPH3A is important for both UBE3A and RAB3A binding we wished to determine whether their binding sites overlap. To identify the critical residues for each interaction we performed site-directed mutagenesis of residues 55 to 72, encompassing about two-thirds of α-helix a1. To minimize the chance that the mutations would affect the α-helical structure, residues were mutated to an alanine or to a leucine (**Fig. 3a, b**). In addition, two charge changes were introduced at position 60 (Arg>Glu) and 65 (Glu>Lys). Mutants were then tested for interaction with UBE3A and RAB3A^Q81L^ in an Y2H assay. As summarized in **Figure 3a (**see also **Supplementary Fig. S4**), the RPH3A mutants V57A, A61L, E65K, E68A, R71A and I72A showed a partial or complete loss of interaction with both UBE3A and RAB3A^Q81L^. The mutants R56A, R60A, R60E and E65A all showed a strong reduction (or complete loss) of UBE3A interaction. For these latter constructs the strength of interaction with RAB3A were essentially unchanged indicating that the observed loss of interaction with UBE3A is specific and not caused by changes in expression levels of the mutant proteins. Notably, based on structural analysis of the RPH3A/RAB3A^Q81L^ complex 9 residues in the RPH3A *α*-helix-1 have been implicated in the RAB3A-RPH3A interaction: E50, I54, V57, I58, R60, A61, M64, E68 and R71 ^17^. Of these 9 residues we tested 7 in our yeast two-hybrid assay and confirmed for 5 of them, V57, A61, E65, E68 and R71, that they are indeed important for RAB3A binding. These results demonstrate the validity of the Y2H technique to study the effect of single amino acid changes on protein-protein interactions and indicate that the binding sites of UBE3A and RAB3A^Q81L^ on RPH3A partially overlap.

**Fig 3.**
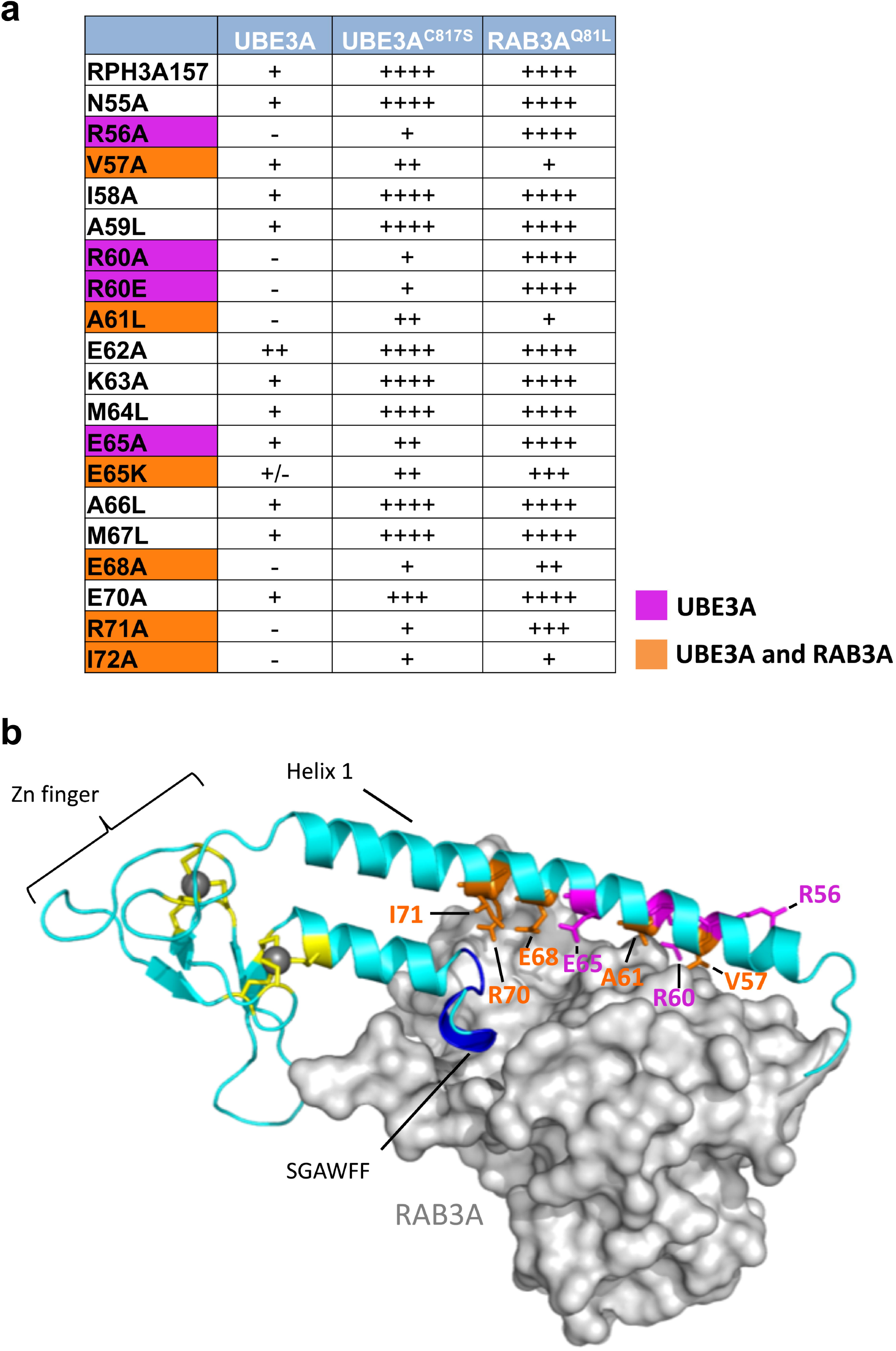
The binding sites of UBE3A and RAB3A on RPH3A partially overlap. **a**) Effect of mutations in the α-helix a1 of RPH3A^1-157^ on the Y2H interaction with UBE3A (WT and C817S mutant) and RAB3A^Q81L^. Interaction strength for each RHP3A mutant is indicated (for details see legend to Figure 1). Residues specific for UBE3A interaction are indicated in magenta, while residues involved in both UBE3A and RAB3A binding are indicated in orange. Scans of the selection plates are presented in **Supplementary Figure S4. b**) Representation of the crystal structure of RPH3A (cyan)-RAB3A (grey) complex ^17^ showing the location of the residues that upon mutation affect the interaction with UBE3A and/or RAB3A. Color-coding is the same as in panel a. The SGWAFF motif is depicted in blue and the Cys residues that coordinate the Zn (grey balls) are depicted in yellow.

The above Y2H data suggest that UBE3A and RAB3A^Q81L^ cannot bind to RPH3A at the same time, most likely due to competition of the overlapping binding sites. To further investigate this, we performed a so called “yeast three-hybrid assay” (Y3H), which is a variation of the Y2H assay where, in addition to two proteins fused to respectively the DNA binding domain (BD) and activation domain (AD), a third (non-fused) protein is expressed. This type of assay allows one to address if a protein interferes with or bridges the interaction between two other (DB- and AD-fused) proteins. We tested the ability of RAB3A^Q81L^ to interfere with UBE3A-RPH3A interaction in a Y3H set-up. UBE3A^C817S^ was used in the assay, as its strength of interaction with RPH3A is comparable to that of RAB3A with RPH3A in our Y2H. As a control, RPH3A^SGAAAA^ mutant was used that is unable to interact with RAB3A^Q81L^ but preserves the ability to interact with UBE3A. As shown in **Figure 4a**, expression of RAB3A^Q81L^ significantly reduced the UBE3A-RPH3A interaction. The effect of RAB3A^Q81L^ on the UBE3A-RPH3A interaction depends on its interaction with RPH3A, as the binding between UBE3A and RPH3A^SGAAAA^, a RPH3A mutant that is unable to interact with RAB3A^Q81L^, was not reduced upon overexpression of RAB3A^Q81L^. Furthermore, overexpression of wild type UBE3A did not reduce the RPH3A-RAB3A^Q81L^ interaction, suggesting that UBE3A is not able to displace the RAB3A^Q81L^ bound to RPH3A (**Fig. 4b**). Finally, we show that there is no interaction between UBE3A and RAB3A^Q81L^, and that RPH3A cannot bridge the interaction between RAB3A^Q81L^ and UBE3A in an Y3H assay (**Fig. 4c**). Together, these data strongly suggest that the GTP loaded form of RAB3A and UBE3A cannot bind RPH3A at the same time.

**Fig 4.**
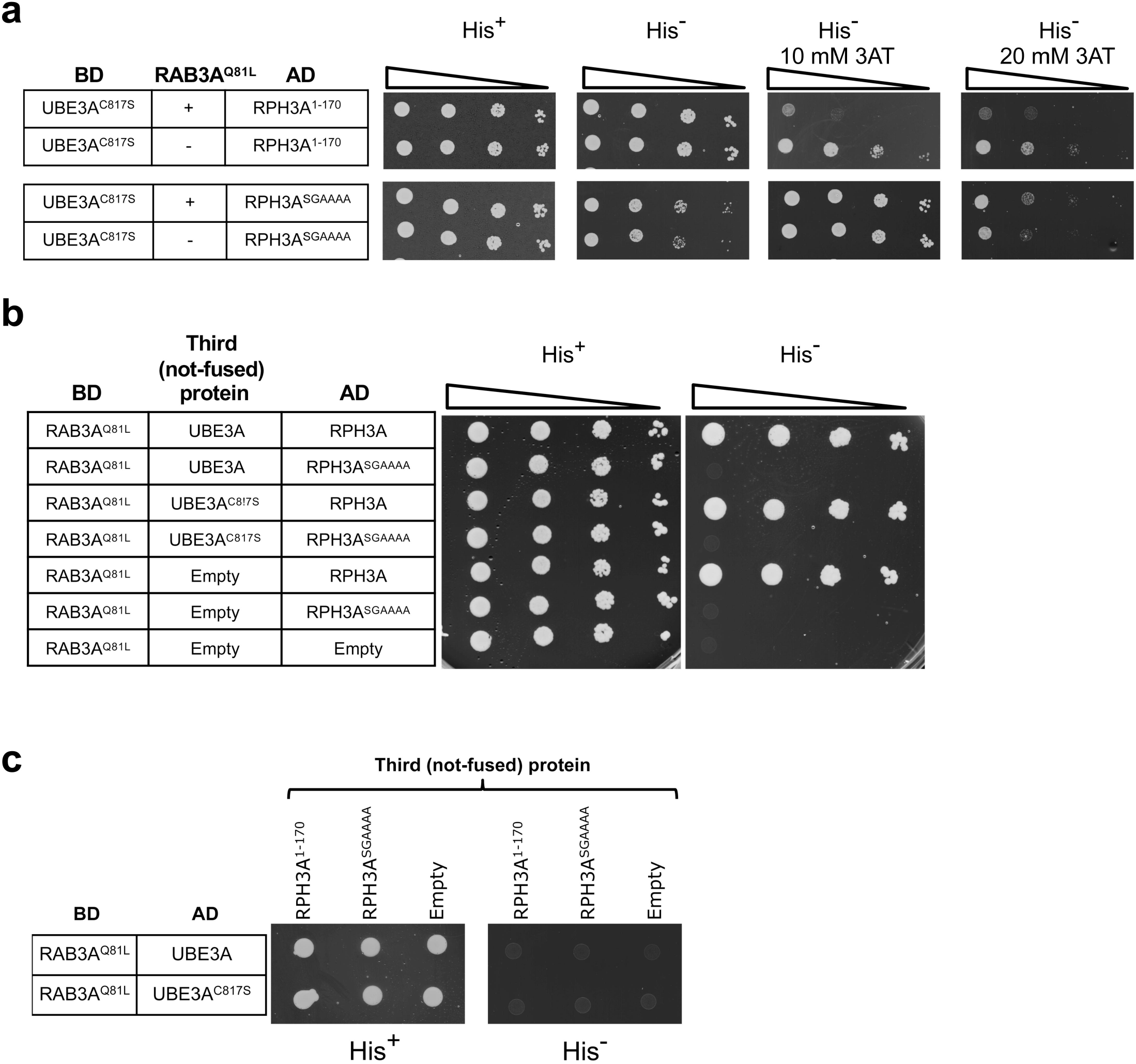
UBE3A and RAB3A cannot simultaneously bind to RPH3A. **a**) RAB3A interferes with the UBE3A-RPH3A interaction. *S. cerevisiae* strain PJ69a was co-transformed with UBE3A^C817S^ (bait), RPH3A^1-170^ or RPH3A^SGAAAA^ (preys), and non-fused RAB3A^Q81L^ or empty vector (pRA1). Cells were spotted on selective plates as indicated. **b**) UBE3A does not interfere with the RAB3A-RPH3A interaction. *S. cerevisiae* strain PJ69a was co-transformed with RAB3A^Q81L^ (bait), RPH3A^1-170^, RPH3A^SGAAAA^ or empty vectors (preys) and non-fused UBE3A, UBE3A^C817S^ or empty vector. Cells were spotted on selective plates as indicated. **c**) RPH3A cannot bridge the RAB3A-UBE3A interaction. *S. cerevisiae* strain PJ69a was co-transformed with RAB3A^Q81L^ (bait), UBE3A or UBE3A^C817S^ (preys), and the indicated non-fused RPH3A constructs or empty vector (pRA1). Cells were spotted on selective plates as indicated.

### Reduced RPH3A protein levels in RAB3A-KO mice are not mediated by UBE3A

Previous studies showed that RPH3A protein levels are significantly decreased in *Rab3a* knock out (KO) mice, while the RNA levels appeared to be unchanged ^14^. This observation strongly suggests that RPH3A protein stability is dependent on the interaction with RAB3A. Given that the binding sites of RAB3A and UBE3A on RPH3A overlap, one plausible explanation for these observations is that in absence of RAB3A, *i*.*e*. in the *Rab3a* KO mice, UBE3A can interact with RPH3A resulting in its ubiquitination and degradation. To test this hypothesis, we deleted UBE3A in *Rab3a* KO mice and determined RPH3A levels in the cortex of adult mice by western blotting. Three different mutants obtained by a cross between AS mice (*Ube3a*^*E113X* 22^) and *Rab3a* KO mice (Jackson laboratories, see material and methods) were used for this analysis: the single gene mutants *Ube3a*^−*/+*^*/RAB3A*^*+/+*^ and *Ube3a*^*+/+*^*/Rab3a*^*-/-*^ and the double mutant *Ube3a* ^−*/+*^*/Rab3a*^*-/-*^. As previously reported^14^, we found strongly reduced RPH3A protein levels in mice lacking both copies of the RAB3A gene (*Ube3a*^*+/+*^*/Rab3a*^*-/-*^) (**Fig. 5**). To investigate if these reduced levels of RPH3A are mediated by UBE3A we analyzed the lysates of mice in which both UBE3A and RAB3A are absent (*Ube3a*^*-/+*^*/Rab3a*^*-/-*^). We found that RPH3A levels in mice lacking both UBE3A and RAB3A (*Ube3a*^*-/+*^*/Rab3a*^*-/-*^) were comparable to those observed in *Ube3a*^*+/+*^*/Rab3a*^*-/-*^ mice, indicating that UBE3A does not mediate the reduction in RPH3A protein levels in *Rab3a* KO mice.

**Fig 5.**
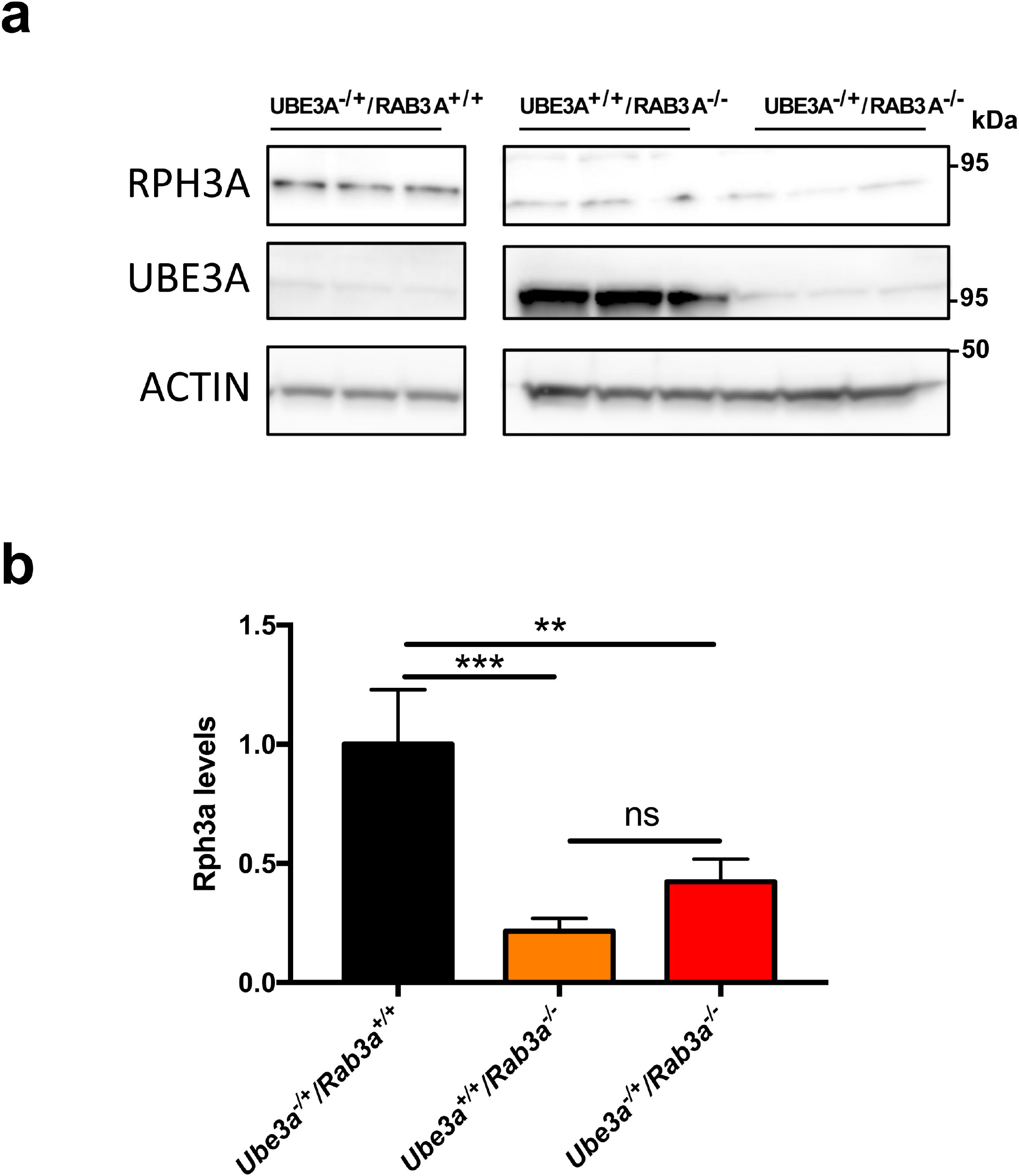
UBE3A is not mediating RPH3A breakdown in *Rab3a-KO* mice. a) *Ube3a*^− *+*^*/Rab3a*^*+/+*^, *Ube3a*^*+/+*^*/Rab3a*^−/–^ and *Ube3a*^−*/+*^*/Rab3a*^−/–^ mice cortexes were lysed and equal amounts of proteins were analysed by SDS-PAGE and RPH3A or UBE3A immunoblotting. ACTIN was used as loading control. **b)** Quantification of RPH3A levels in *Ube3a*^− *+*^*/Rab3a*^*+/+*^, *Ube3a*^*+/+*^*/Rab3a*^−/–^ and *Ube3a*^−*/+*^*/Rab3a*^−/–^ mice cortexes (one way ANOVA, multiple comparisons, *Ube3a*^−*/+*^*/Rab3a*^*+/+*^ n=3, *Ube3a*^*+/+*^*/Rab3a*^−/–^ n=3, *Ube3a*^−*/+*^*/Rab3a*^−/–^ n=3, error bars depict +/- SD). Note: all data presented in this figure come from the same gel (see **Supplementary Figure S9** for original blots used for the quantification).

### RPH3A is ubiquitinated *in vivo* in an UBE3A-dependent manner

While the above mouse experiments strongly suggest that UBE3A is not involved in the regulation of RPH3A levels, these results do not exclude the possibility that RPH3A is a *bona fide* ubiquitination target of UBE3A. To determine if RPH3A is ubiquitinated *in vivo* in an UBE3A-dependent manner we performed diGly mass spectrometry on pooled cortical tissue obtained from seven P28 *Ube3a* ^−/–^ mice and seven wild type littermates (^23^; **Fig. 6a, b and c**). We found that RPH3A is ubiquitinated at three different sites in wild type mouse brain while in *Ube3a*^−/–^ mice the ubiquitinated RPH3A peptides were either absent or strongly reduced (**Fig. 6a**). Importantly, total intensities of proteins and diGly peptides were comparable between wild-type and mutant samples (**Fig. 6b, c**). These results unambiguously show that RPH3A is ubiquitinated *in vivo* and strongly suggest that RPH3A ubiquitination is dependent on the presence of UBE3A. Notably, total RPH3A proteins levels were comparable between wild type mice and *Ube3a* ^−/–^ mice (**Supplementary Fig. S5**). To investigate if RPH3A is a direct ubiquitination target of UBE3A, we modified a previously described bacterial ubiquitination system ^24^. With this system, it can be determined whether a protein is ubiquitinated in the presence of a *single* E3 ligase and the appropriate E2 in the context of a (bacterial) cell. We first confirmed the validity of the system by co-transforming *E. coli* cells with UBE3A (WT or C817S), UBCH7 (the cognate E2 for UBE3A), UBA1 (rabbit E1 enzyme), ubiquitin and RING1B, a well-established target of UBE3A^25^. In the presence of active UBE3A and all components of the ubiquitination cascade, a smear of slower migrating bands was observed for RING1B (**Fig. 6d**), indicative of poly-ubiquitin chain formation. Ubiquitination of RING1B was not observed when the catalytically inactive UBE3A (UBE3A^C817S^) was present or in the absence of one of the components of the ubiquitination cascade. Target selection by UBE3A is highly specific in this assay as UBE3A failed to ubiquitinate the control protein luciferase. Importantly, we show that ARC, a protein that was suggested to be a substrate of UBE3A^26^ but was subsequently shown to be regulated at the transcriptional level^27^ is not ubiquitinated by UBE3A (**Fig. 6e**). Since it was previously reported that some E3 ligases are promiscuous with respect to their substrate specificity in *in vitro* ubiquitination assays using purified proteins (*e*.*g*. the RING type E3 ligase MDM2^27^), we tested MDM2 as well as NEDD4 (a HECT type E3 ligase) for their ability to ubiquitinate RING1B in the bacterial ubiquitination assay. We found that neither E3 ligase ubiquitinated the UBE3A substrate RING1B, further highlighting the specificity of the system (**Fig. 6f**). Having validated the specificity and selectivity of the bacterial ubiquitination assay, we next asked the question if RPH3A is ubiquitinated by UBE3A. Interestingly, when RPH3A was co-expressed with UBE3A and all components of the ubiquitination cascade we observed one predominant slower migrating RPH3A band indicative of mono- or multi-mono-ubiquitination (**Fig. 6g**).

**Fig 6.**
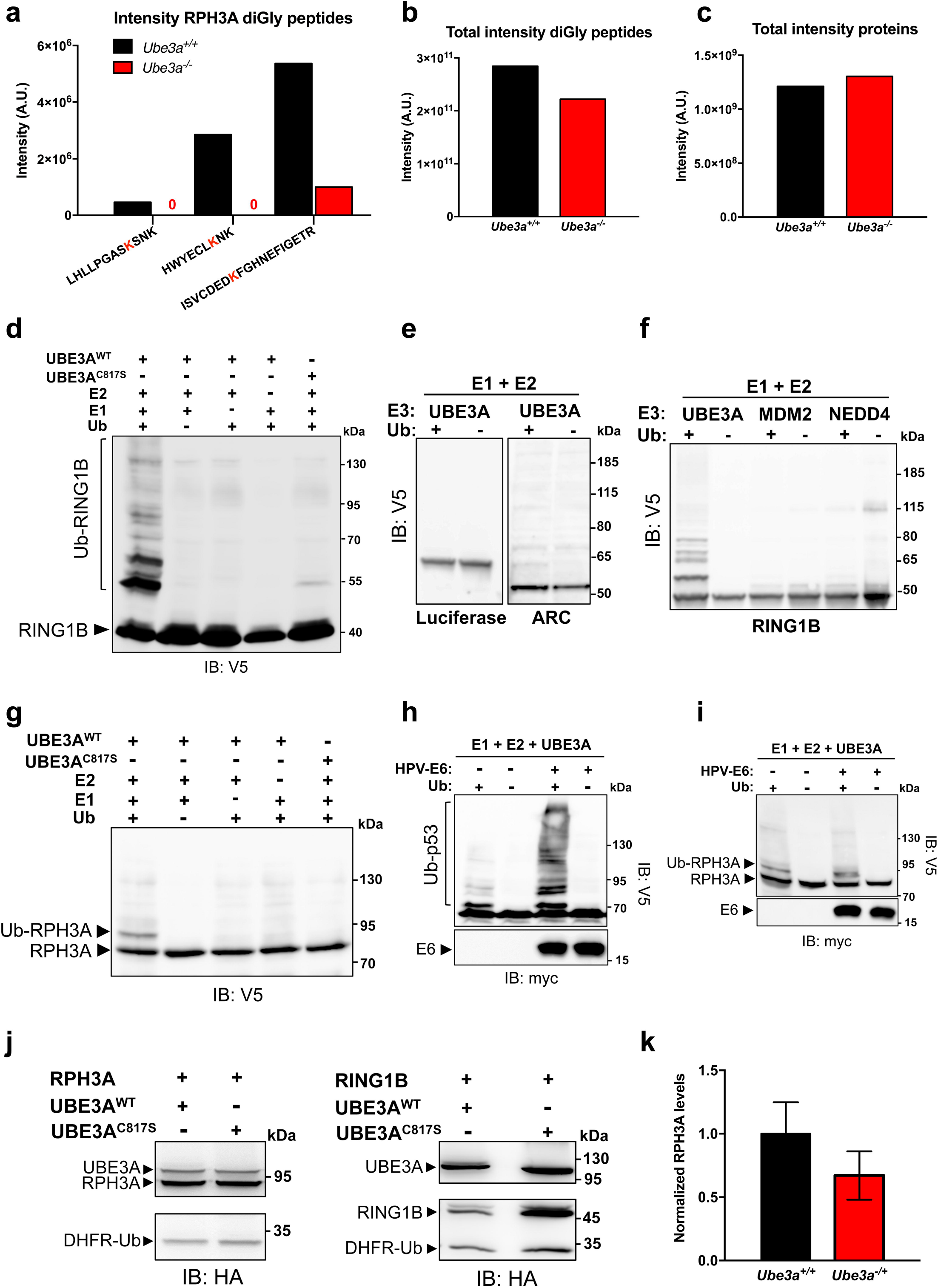
RPH3A ubiquitination *in vivo* is UBE3A-dependent and does not affect RPH3A stability. **a**) Intensities of three unique RPH3A diGly peptides in pooled cortical tissue obtained from seven P28 *Ube3a*^m–*/p-*^ (*Ube3a* KO) mice and seven wild type (*Ube3a*^m+*/p+*^) littermates. After digestion of protein lysates with trypsin, diGly peptides were enriched by immunoprecipitation with ubiquitin remnants motif (K-ε-GG) antibodies and identified by nanoLC-MS. **b**) Total intensity of diGly peptides in *Ube3a*^m+*/p+*^ and *Ube3a*^m–*/p-*^ brain lysates. **c**) Total intensity of proteins in *Ube3a*^m+*/p+*^ and *Ube3a*^m-*/p-*^ brain lysates. **d**. Ubiquitination of the positive control RING1B^I53S^ (a catalytically inactive mutant of the E3 ligase RING1B) in the bacterial ubiquitination assay. *E. coli* cells co-expressing the indicated combination of HA-UBE3A, HA-UBE3A^C817S^, V5-RING1B^I53S^, UbcH7, E1 and ubiquitin (Ub) were lysed and equal amounts of total lysates were analyzed by SDS-PAGE and anti-V5 immunoblotting. Note that RING1B ubiquitination is only observed in the presence of wild type UBE3A and all components of the ubiquitination cascade, but not when the catalytically dead UBE3A mutant (UBE3A^C817S^) is present (**e**). Analysis of luciferase and ARC in the bacterial ubiquitination assay. V5-Luciferase or V5-ARC were co-expressed in bacteria with HA-UBE3A, E1, E2 and either with (+) or without (-) ubiquitin and total lysates were analyzed by immunoblotting. **f**) MDM2 and NEDD4 do not ubiquitinate the UBE3A target RING1B in bacteria. HA-UBE3A, HA-MDM2 or HA-NEDD4 were co-expressed in bacteria with V5-RING1B^I53S^ and the indicated components of the ubiquitination cascade and total lysates were analyzed immunoblotting. **h)** *E. coli* cells co-expressing p53 and the indicated combination of HA-UBE3A, myc-E6 (HPV16) and ubiquitin (Ub) were lysed and total lysates were analyzed by immunoblotting using the indicated antibodies. **i)** *E. coli* cells co-expressing RPH3A and the indicated combination of HA-UBE3A, myc-E6 (HPV16) and ubiquitin (Ub) were lysed and total lysates were analyzed by immunoblotting. **j**) HEK293T-UBE3A^KD^ cells were co-transfected with HA-UBE3A or HA-UBE3A^C817S^ and DHFR-HA-ubiquitin-fused HA-RPH3A or HA-RING1B^I53S^. The levels of DHFR-HA-Ub served as an internal control to determine the effect of UBE3A on the levels of RPH3A and RING1B^I53S^. For details, see text. RING1B^I53S^, a known target of UBE3A, was used as positive control. Total protein extracts were prepared and analyzed by SDS-PAGE and anti-HA immunoblotting. Full-length blots for **Figs. 6a-d** are presented in **Supplementary Figure S10. k**) Quantification of RPH3A in cortical lysates of WT (*Ube3a*^m+*/p+*^) and AS (*Ube3a*^m-*/p+*^) mice (t-test, WT n=3, AS n=3, error bars depict +/- SEM). Blot used for quantification is presented in **Supplementary Figure S6** and full-length blots are presented in **Supplementary Figure S11**.

The human papilloma virus (HPV) E6 protein has been reported not only to mediate UBE3A-dependent degradation of the tumor suppressor protein p53 ^2^, but also to act as an allosteric activator of UBE3A, enhancing both UBE3A self-ubiquitination as well as target ubiquitination ^28,29^. To test whether UBE3A is able to poly-ubiquitinate RPH3A in the presence of the HPV E6 activator we performed ubiquitination assays in bacteria that also express the viral E6 protein (**Fig. 6h and i**). Co-expression of the HPV E6 protein did not change the UBE3A-mediated ubiquitination pattern on RPH3A compared to cells not expressing E6 (**Fig. 6i**), while p53 ubiquitination was substantially increased, both in the amount and length of the ubiquitin chains, in cells that express E6 (**Fig. 6h**). Together, these results show that RPH3A is a direct, E6-independent, ubiquitination substrate of UBE3A.

### UBE3A-dependent ubiquitination of RPH3A does not serve as a degradation signal

Our results in the bacterial ubiquitination assay suggest that the modification of RPH3A by UBE3A is non-canonical, *i*.*e*. mono or multi-mono ubiquitination rather than poly-ubiquitination. While poly-ubiquitination (e.g. K48-linked ubiquitin chains) is often associated with proteasome-mediated degradation, mono-ubiquitination/multi-mono-ubiquitination is usually involved in protein trafficking and signaling ^4^. To determine if this type of ubiquitination can result in degradation of RPH3A in mammalian cells we used the well-characterized dihydrofolate reductase (DHFR)-ubiquitin fusion protein system ^27^. In this system the protein of interest is fused C-terminally to DHFR-ubiquitin, resulting in the expression of a single polypeptide that is co-translationally and quantitatively cleaved by a ubiquitin-specific protease into two separate proteins: the protein of interest and DHFR-ubiquitin. By comparing the relative levels of the protein of interest and DHFR-ubiquitin, the effect of UBE3A on the stability of the target can be directly quantified. HEK293T-UBE3A^KD^ cells in which UBE3A is partially knocked down were co-transfected with RPH3A or RING1B fused to DHFR-Ub, and either wild type UBE3A or the catalytically inactive UBE3A^C817S^ mutant. 48 hours post-transfection total protein lysates were prepared and analyzed by immunoblotting (**Fig. 6j**). Comparable RPH3A levels were observed in cells co-transfected with either UBE3A or UBE3A^C817S^, while RING1B levels were significantly decreased by co-expression of UBE3A, but not by co-expression of UBE3A^C817S^. Finally, we tested if UBE3A affects RPH3A levels *in vivo* by performing immunoblot analysis on mouse brains derived from AS (*Ube3a*^m–/p+^) and WT mice. As shown in **Figure 6k** (and **Supplementary Fig. S6)**, we observed no significant differences in the RPH3A levels between AS and WT mice by immunoblot analysis. These results are in line with the proteomics data that showed comparable RPH3A proteins levels in cortical lysates of wild type mice and *Ube3a*^−/–^ mice (**Supplementary Fig. S5**). Taken together, these data demonstrate that while RPH3A is ubiquitinated by UBE3A within cells, in the mouse brain RPH3A is not targeted by UBE3A for (proteasomal) degradation. These results are consistent with our ubiquitination data that show mono-/multi-monoubiquitination of RPH3A by UBE3A, the exact function of which remains to be determined.

## Discussion

In this study we have identified the pre-synaptic protein RPH3A as a novel binding partner of UBE3A. Deletion analysis showed that an N-terminal fragment of RPH3A (residues 39-87) is necessary and sufficient to bind UBE3A. At the core of this RPH3A fragment is the α-helix a1 of RPH3A (residues 48 to 85), a region that is also involved in RAB3A binding. Alanine-scanning mutagenesis of this region revealed that the binding sites of UBE3A and RAB3A on RPH3A partially overlap, suggesting a competing binding mechanism. Consistent with this, we show in an Y3H set-up that RAB3A interferes with the UBE3A-RPH3A interaction. While this work was in progress, Martinez-Noël and colleagues also found RPH3A as an interaction partner of UBE3A in an Y3H screen with the catalytically inactive UBE3A and in the presence of the HPV E6 protein, but not when the catalytically active UBE3A was used or in the absence of E6 ^30^. By contrast, our interaction data indicates that the catalytically active UBE3A can bind RPH3A in the absence of E6, although the interaction is relatively weak and a stronger interaction is observed with the catalytically inactive UBE3A. Therefore, our data indicate that the UBE3A-RPH3A interaction does not require the presence of the viral E6 protein. In further support of this we recently identified the region in UBE3A required for RPH3A binding and showed that the RPH3A binding domain is located between residues 253 and 274 of UBE3A^19^, a region that is distant from the E6 binding site^31^. We now show that an AS-linked mutation in the RPH3A binding region (p.Leu273Phe) reduced the interaction with UBE3A. Collectively, our results indicate that RPH3A is a *bona fide* interaction partner of UBE3A.

We investigated whether RPH3A is not only a binding partner but also a ubiquitination substrate of UBE3A. We show that RPH3A is ubiquitinated in an UBE3A-dependent manner in mouse brain and is a direct target of UBE3A in a bacterial ubiquitination assay that we adapted for UBE3A target analysis. Although we observed poly-ubiquitination of well-established targets, we found that RPH3A is (multi) mono-ubiquitinated in this assay. Notably, while E6 enhances the UBE3A-RPH3A interaction (^30^and **Supplementary Fig. S7**) we did not observe a change in the level or the length of the ubiquitin modification on RPH3A in the presence of E6. Consistent with a non-degradative role of the UBE3A-mediated RPH3A modification we show that RPH3A levels are not affected by overexpression of UBE3A in HEK293T cells. We also investigated the *in vivo* effect of UBE3A on RPH3A levels. We compared the RPH3A levels in AS mice (*Ube3a*^*–/+*^) to that of wild type mice (*Ube3a*^*+/+*^) by immunoblotting and proteomics and found no difference (**Fig. 6k and Supplementary Fig. S5**). We also made use of *Rab3a* KO mouse mutants, for which it had been previously shown that this effects RPH3A protein (but not RPH3A RNA) levels^14^, suggesting enhanced breakdown of RPH3A in the absence of RAB3A. We confirmed this finding, but despite the overlapping UBE3A and RAB3A binding sites, the extreme low levels of RPH3A could not be rescued by also deleting *Ube3a*. This indicates that in the absence of RAB3A, RPH3A is not targeted by UBE3A for degradation. In conclusion, our data show that the UBE3A-RPH3A interaction leads to mono/multi-monoubiquitination of RPH3A, a modification that does not affect the stability of the protein neither in cells nor in the mouse brain. Mono- and multi-mono-ubiquitination have been previously described as post-translational modifications involved in chromatin remodeling and DNA repair, and mono-ubiquitination often has a role in protein signaling ^4^. In light of this, we hypothesize that UBE3A-dependent ubiquitination of RPH3A may regulate the activity of the protein, perhaps by determining the amount of protein capable of RAB3A interaction. Further experiments will be needed to investigate this.

One of the first studies on RPH3A described its role in the regulation of exocytosis^32^. Burns and colleagues revealed how injection of the full-length RPH3A protein into giant squid synapses inhibits neurotransmitter release by preventing the priming of synaptic vesicles. This phenomenon is most likely due to the continuous binding of excessive RPH3A to RAB3A, which inhibits GTP hydrolysis by RAB3A thereby attenuating the vesicle cycle. On the other hand, the same study showed that synaptic injection of RPH3A fragments, either the N-terminal domain involved in RAB3A binding or the C-terminal C2 domains, results in the accumulation of un-docked vesicles. In this case RPH3A fragments are thought to act as dominant negative regulators towards potential binding partners of RPH3A involved in the docking phase. Since UBE3A and RAB3A compete for RPH3A binding through interaction with the N-terminal α-helix, a plausible hypothesis would be that ubiquitination of RPH3A by UBE3A interferes with RPH3A-RAB3A interaction, and thereby has an effect on the number of docked vesicles at the pre-synaptic membrane. Further experiments will be needed to determine whether the absence of UBE3A in AS individuals has an effect on vesicles docking and neurotransmitter release.

RPH3A has also been implicated to play a role in endocytosis at the synaptic bouton^33,34^. Higher levels of RPH3A contribute to an increased number of clathrin-coated vesicles (recycled vesicles). Since RPH3A is not associated with clathrin-coated vesicles ^14,35^ it has been suggested that the effector plays a role at an early stage of the endocytic pathway. In the model proposed by Coppola *et al*. ^33^, immediately after the dissociation from RAB3A, RPH3A is free to bind, through its N-terminus, RABAPTIN5, a known component of the endocytic machinery ^36^. This interaction is required to initialize vesicles endocytosis leading to the assembly of the required multiprotein complex. This last aspect is particularly interesting for the Angelman Syndrome pathophysiology since Judson *et al*. reported that AS neurons show an impaired endocytic machinery ^37^.

Although RPH3A is known as a protein involved in the regulation of exo- and endocytosis processes at presynaptic sites, a recent study showed that RPH3A is also present in dendritic spines ^38^. This study indicated that at post-synaptic spines RPH3A acts as inhibitor of endocytosis, and by forming a ternary complex with GluN2A (a subunit of the NMDA receptor) and PSD-95, RPH3A is able to stabilize the NMDA receptor surface expression^38^. Different studies have reported that GluN2A expression throughout the brain is developmentally regulated with its levels in mouse brain sharply increasing from the second postnatal week ^39,40^. If this finely tuned process is disturbed through mis-regulaton of RPH3A ubiquitination by UBE3A, it could potentially affect neuronal function in AS, but this has to be further investigated.

UBE3A immuno-labelling has been shown to be present in the pre- and post-synaptic compartment of the synapse ^41-43^, and several electrophysiological studies showed impaired synaptic function in AS mice ^10,26,44,45^. Thus, in combination with the aforementioned pre and postsynaptic role of RPH3A and our findings that RPH3A is a *bona fide* substrate of UBE3A, the regulation of RPH3A by UBE3A could point towards an important function. However, it remains to be determined to what extent these findings are relevant to the pathophysiology of Angelman Syndrome. In particular, we recently showed that the changes in synaptic function of AS mice are also observed if only the nuclear UBE3A isoform is deleted ^19^. This finding could suggest that the core phenotypes of AS are independent of the UBE3A-RPH3A interaction. Thus, more research is needed to address the clinical significance of our findings.

## Material and methods

### Plasmid construction

UBE3A constructs were amplified from mouse cDNA using a combination of forward and reverse primers as given in **Supplementary Table S1**, thereby introducing restriction sites at the 5’ and 3’ end of the gene fragment. PCR products were cloned by A-tailing into pGEMTeasy (Promega) and sequenced. RPH3A constructs were amplified from Image Clone BC053519 using the primers indicated in **Supplementary Table S1**. Single amino acid changes in RPH3A, RAB3A and UBE3A were introduced by site-directed mutagenesis following the manufacturer’s protocol (Agilent). pCI-HA-NEDD4 (#27002), pGEX-4T-MDM2-WT (#16237) and pCMV-IRES-Renilla Luciferase-IRES-Gateway-Firefly Luciferase (pIRIGF) (#101139) expression constructs were purchased from Addgene. For Y2H, RPH3A constructs were cloned AscI-NotI into the prey vector pGADT7mod, which is a derivative of pGADT7 (Clontech) containing a modified multiple cloning site. For mammalian expression, RPH3A constructs were cloned AscI-NotI into a pEGFPn3 (Clontech) derived plasmid deprived of the GFP (pRA196) and BamHI-NotI into the DHFR-ubiquitin fusion plasmid (see below) cut with BamHI and PspOMI (pBD242). UBE3A constructs were cloned AscI-NotI into pYR22, a single copy bait vector for yeast two-hybrid assays. For constitutive expression of UBE3A, RAB3A and RPH3A constructs in yeast three-hybrid assays the ORFs were cloned Asc/Not in the previously described yeast expression vector pRA1^46^. For the bacterial ubiquitination system^24^ and co-immunoprecipitation experiments we designed novel compatible expression vectors. pMB419 is a derivative of pCOLA-Duet1 (Novagen) that encodes a single T7 promoter followed by a *lac* operator, a ribosome binding site (*rbs*), a V5 epitope tag and a multiple cloning site (MCS). The vector has the ColA1 origin of replication and the ampicillin resistance gene. For bacterial ubiquitination experiments RPH3A^FL^ was cloned AscI-NotI in pMB419. pYW5 is a polycistronic expression vector derived from pRSF-Duet1 (Novagen). The vector encodes a single T7 promoter followed by a *lac* operator, a rbs, a HA epitope tag, a multiple cloning site (MCS), a second *rbs*, a myc epitope and a second MCS. The vector carries the RSF origin of replication and the Spectinomycin/Streptomycin resistance gene derived from pCDF-Duet1 (Novagen). UBE3A constructs (wild type and catalytically inactive mutant) were cloned AscI-NotI in the first MCS of pYW5. pCI-HA-NEDD4 was digested KpnI-NotI and the NEDD4 insert was cloned into AscI-NotI digested pYW5 in a three-fragment ligation using a double stranded oligonucleotide with AscI-KpnI overhangs. MDM2 (*H. sapiens* wild type gene) was PCR amplified from pGEX-4T-MDM2-WT, thereby introducing a 5’ AscI and 3’ NotI site. The resulting MDM2 PCR product was digested AscI-NotI and cloned into AscI-Not cut pYW5. Four derivatives of the generic plasmid pGEN4 (generous gift of Gali Prag), which co-expresses His6-tagged ubiquitin, yeast Ubc4 and rabbit E1, were constructed. In pMB277 the His6-tagged ubiquitin was replaced by wild type (untagged) ubiquitin and yeast Ubc4 by mouse UbcH7. To construct pMB278 the gene encoding wild type ubiquitin was deleted from pMB277. Deletion of the E2 in pMB277 resulted in pMB316. Similarly, deletion of the E1 in pMB277 gave rise to pMB318. The RING1B^I53S^ construct in the dehydrofolate reductase (DHFR)-ubiquitin fusion plasmid was a generous gift of Martin Scheffner^27^. For bacterial ubiquitination experiments targets were cloned in the pMB419 MCS as follows. The RING1B-I53S fragment isolated from the DHFR-ubiquitin fusion plasmid was cloned BamHI-SalI, Firefly Luciferase was PCR amplified from the original construct, thereby introducing a 5’ AscI and 3’ NotI site, and cloned in AscI-NotI, and the Arc containing plasmid pRA32^46^ was digested AscI-NotI and cloned similarly in pMB419. For co-immunoprecipitation experiments a second pRSF-Duet1 derivative was designed. Plasmid pMB280 is a polycistronic expression vector similar to pYW5, except that it has a V5 epitope tag preceding the first MCS and a HA epitope tag preceding the second MCS. RPH3A constructs were cloned BamHI-SalI in the first MCS, generating V5-tagged proteins, while UBE3A (or GFP) was inserted in the second MCS using AscI and NotI restriction sites, introducing a HA tag at the N-terminus of UBE3A (or GFP). For some of the bacterial ubiquitination experiments pMB280 was modified to express a third gene by insertion of a double-stranded (ds)-oligonucleotide between the NotI and AvrII sites. The ds-oligonucleotide encodes a *rbs* followed by a Myc tag and a MCS. The resulting plasmid pMB323 can express three different proteins each with a different epitope tag (HA, V5 or Myc) from a single T7 promoter. The HPV E6-type 16 the gene was amplified from plasmid #65 (a generous gift of Jon A Huibregtse) introducing a 5’ SbfI- and a 3’ PacI site, and cloned in the third MCS of pMB323, giving rise to pMB336. pMB336 was used to clone UBE3A (wild type and catalytically inactive mutant) in the second MCS and the targets (RPH3A or p53 [generous gift of Jon Huibregtse]) in the first MCS. A complete list of the plasmids employed in this study can be found in **Supplementary Table S2**. All constructed plasmids were sequence verified with Sanger sequencing (Macrogen Europe). Details of the construction of these plasmids and their full sequences are available upon request.

### Yeast two-hybrid assay (Y2H)

*S. cerevisiae* strains used for yeast two-hybrid analysis were Y187 (*MATα, ura3-52, his3-200, ade2-101, trp1-901, leu2-3, 112, gal4Δ, met*^*–*^, *gal80Δ, URA3::GAL1*_*UAS*_*-GAL1*_*TATA*_*-lacZ*; Clontech) and Y2H Gold (*MATa, ura3-52, his3-200, ade2-101, trp1-901, leu2-3, 112, gal4Δ, gal80Δ, met*^*–*^, *LYS2 :: GAL1*_*UAS*_*-Gal1*_*TATA*_*-His3,GAL2*_*UAS*_*-Gal2*_*TATA*_*-Ade2, URA3* : : *MEL1*_*UAS*_*-Mel1*_*TATA*_*-AUR1-C MEL1*; Clontech). Cells were grown at 28 °C in rich medium or in minimal glucose medium as previously described ^47^.

Y2H interactions between bait constructs (mouse UBE3A or mouse RAB3A^Q81L^) and prey constructs (mouse RPH3A) were assessed using a mating protocol, as previously described ^48^.

### Yeast three-hybrid assay (Y3H)

For the yeast three-hybrid analysis *S. cerevisiae* strain PJ69-4a (*MAT*a *trp1-901 leu2-3,112 ura3-52 his3-200 gal4Δ gal80Δ lys2::GAL1-HIS3 GAL2-ADE2 met2::GAL7-lacZ*) was transformed with the bait constructs (UBE3A or RAB3A), the prey constructs (RPH3A) and a third plasmid constitutively expressing either RAB3A or RPH3A. Transformants were then selected on triple selective agar plates (lacking leucine, tryptophan and uracil) and spotted as previously described ^47^.

### Bacterial ubiquitination assay

*E. coli* BL21-GOLD (DE3) [B F^−^ *ompT hsdS*(r_b-_ m_b-_) *dcm*^*+*^ *Tet*^*r*^ *gal λ(DE3) endA Hte*] cells were co-transformed with bacterial expression constructs and selected on LB agar (1% (w/v) Bacto tryptone, 0.5% (w/v) Bacto yeast extract, 1% (w/v) NaCl, 1.5% agar) containing half the concentration of antibiotics as needed (ampicillin, 25 μg/ml; kanamycin, 15 μg/ml; streptomycin/spectinomycin, 25 μg/ml). Transformants were inoculated in LB medium supplemented with 2% glucose, 50 mM Tris (pH 8.0) and antibiotics as needed, and grown ON at 37 ^°^C while shaking (200 rpm). The next morning cells were inoculated at an OD_600_ of 0.2 and grown at 21°C to an OD_600_ of 0.7. Cells were then induced by the addition of 0.5 mM Isopropyl B-D-1-thiogalactopyranoside (IPTG, INVITROGEN Life Technologies) and further grown at 21°C for 16-18 h. Bacterial pellets of 20 OD_600_ units were lysed in 0.5 ml cold lysis buffer (50 mM Sodium-phosphate pH 8.0, 300 mM NaCl, 5% glycerol, 5 mM 2-Mercaptoethanol, 1 mM PMSF [Sigma]), RNase [0.01 mg/ml], DNase [0.01 mg/ml) with protease inhibitor cocktail (Roche)) by sonication (Sanyo soniprep). Debris were removed by centrifugation (13,000 xg, 30 min) and cleared lysates were used for immuneblot analysis.

### *E.coli* co-immunoprecipitation assay

*E. coli* cells overexpressing the constructs of interest were grown and lysed as described above. 50 µl of the cleared bacterial lysates (corresponding to 2 OD_600_ units) were five times diluted in lysis buffer without DNase and RNase, and containing 0.5% Igepal CA630 (Sigma-Aldrich). Samples were pre-cleared by the adding 25 µl of 50% Sepharose CL-6B beads (Sigma-Aldrich) previously washed with a wash buffer containing 50 mM Na-Pi-buffer pH 8.0, 300 mM NaCl, 5% glycerol and 0.5% Igepal for 30 minutes at 4 ^°^C (while rotating). The beads were pelleted and the supernatants were incubated overnight with 7µg HA antibody (Sigma-Aldrich) or 10µg mouse IgG antibody (Santa Cruz Biotechnology, Inc) at 4^°^C. Protein G Sepharose 4 Fast flow beads (GE Healthcare Life Sciences) washed twice with wash buffer were added to each IP and incubation was continued for 2 hours at 4^°^C. Beads were pelleted, washed three times with wash buffer and once with 50 mM Na-Pi-buffer [pH 8.0] containing 150 mM NaCl, and bound proteins were eluted by incubating the beads for 10 min at 37^°^C with Laemmli sample buffer (62.5 mM Tris-HCl pH 6.8, 2% SDS, 10% glycerol, 0.004% bromophenol blue) supplemented with 50mM DTT. Approximately 40% of the eluates (corresponding to 0.8 OD_600_ units) and ∼10% of the total lysates (corresponding to 0.2 OD_600_ units) were loaded on NuPAGE gels, and analyzed by immuno-blotting.

### Transfection in HEK293T cells

HEK293T cells were grown in DMEM supplemented with 10% foetal calf serum (FCS) and 5% pen strep (PS) at 37°C and 5% CO2. They were transfected with the respective constructs following the JetPRIME transfection system protocol (Polyplus). Total cell lysates were prepared and analysed as previously described ^48^.

### Mice and sample preparation

Mice used for the diGly mass spectrometry experiments (**Figure 6a-c**) were obtained from crosses between previously described *Ube3a*^*m–/p+*^ mice (AS mice, also referred to as *Ube3a*^*–/+*^ in the text) (*Ube3a*^*tm1Alb*^; MGI 2181811)^49^ yielding both *Ube3a*^*m–/p–*^ (*Ube3a* KO) and *Ube3a*^*m+/p+*^ (wild type mice, also referred to as *Ube3a*^*+/+*^) littermates. Cortices taken from 7 P28 *Ube3a*^*m–/p–*^ mice and seven P28 *Ube3a*^*m+/p+*^ controls were separately pooled and taken up in a lysis buffer consisting of 100mM Tris (pH 8.5), 8M urea and 0.15% sodium deoxycholate. Samples were transferred to the in-house proteomics facility.

For the experiment in **Figure 6k**, we used *Ube3a*^*m–/p+*^ mice as previously described ^49^. *Ube3a*^*tm1Alb*^ mice were maintained (>40 generations) in the 129S2 background (full name: 129S2/SvPasCrl) by crossing male *Ube3a*^*m+/p–*^ mice with female 129S2 wild-type mice. For immunoblotting assays, female *Ube3a*^*tm1Alb*^ (*Ube3a*^*m+/p–*^) mice were crossed with male WT mice to obtain both *Ube3a*^*m–/p+*^ and WT mice in the same litter.

For the experiment in **Figure 5**, we used *Ube3a*^*E113X*^ mice (*Ube3a*^*tm2Yelg*^; MGI:5911277), as previously described ^22^, crossed with *Rab3a*^*–/+*^mice. *Ube3a*^*E113X*^ mice were back crossed for 13 generations in B6 background by crossing *Ube3a*^*E113X*^ males with C57BL/6J females. *Rab3b*^*tm1Sud*^ *Rab3a*^*tm1Sud*^ *Rab3d*^*tm1Rja*^ *Rab3c*^*tm1Sud*^ (Jackson lab 006375) were crossed with C57BL/6J mice to obtain *Rab3b*^*–/+*^*Rab3a*^*–/+*^*Rab3d*^*–/+*^*Rab3c*^*–/+*^. *Rab3b*^*–/+*^*Rab3a*^*–/+*^*Rab3d*^*–*^ *+Rab3c*^*–/+*^ mice were subsequently crossed once more with C57BL/6J mice to obtain *Rab3b*^*+/+*^*Rab3a*^*–/+*^*Rab3d*^*+/+*^*Rab3c*^*+/+*^, above referred to as “*Rab3a*^*–/+*^”. *Rab3a*^*–/+*^ mice were maintained for 6 generations in C57BL/6J background by crossing Rab3a^*–/+*^ mice with C57BL/6J. Subsequently, Rab3a^*–/+*^ mice were crossed with *Ube3a*^*E113X*^ mice. From the offspring, male *Ube3a*^*m+/p+*^*/Rab3a*^*+/–*^ were crossed with female *Ube3a*^*m+/p–*^*/Rab3a*^*+/–*^ *mice*, to obtain *Ube3a*^*m–/p+*^/*Rab3a*^*+/+*^, *Ube3a*^*+/+*^*/Rab3a*^*-/-*^ and *Ube3a*^*m–/p+*^/*Rab3a*^*–/–*^ mice used in this study.

Mouse cortex was isolated from adult mice and lysed in 2x Laemmli sample buffer. 20μg of tissue lysates were loaded on SDS-PAG and analysed by UBE3A and ACTIN immunoblotting. All animal experiments were conducted in accordance with the European Commission Council Directive 2010/63/EU (CCD approval AVD101002016791) and approved by the local ethics committee (Instantie voor Dierenwelzijn; IVD).

### diGly mass spectrometry

Ubiquitination sites were determined by a proteomics method based on enrichment of peptides carrying a di-Glycine (diGly) remnant as a result of trypsinization of ubiquitinated proteins, as described previously^23^. The stoichiometry of ubiquitination was determined by relative quantitation of diGly peptides based on their mass spectral intensities.

## Acknowledgements

We are grateful to Gali Prag for the pGEN4 plasmid, to Martin Scheffner for the UBE3A^KD^ cells and the DHFR-Ub plasmid, to Jon Huibregtse for the p53 and E6 plasmids, to Boris Bleijlevens for providing protein modelling advice, to Geeske van Woerden, Diana Rotaru and Minetta Elgersma for the constructive criticism and to Minetta Elgersma and Mehrnoush Aghadavoud Jolfaei for colony management and genotyping. This research was supported by grants from the Netherlands Organization of Scientific Research (NWO-ZoN-MW; 91209046 and 91216045) to BD and YE, and the Angelman Syndrome Alliance (ASA) to BD and YE.

## Author contributions

BD, RAT and YE designed the experiments; RAT, MP, EM, IdG, MB, IZ, JD and YW performed the experiments; BD and RAT wrote the manuscript; BD, YE and RAT reviewed and edited the manuscript; BD and YE supervised the project; BD and YE acquired the funding.

## Declaration of Interests

The authors declare no competing interests

## Data Availability

No datasets were generated or analyzed during the current study

## Notes

### Competing Interest Statement

The authors have declared no competing interest.

