## Supplemental data for "Mono-ubiquitination of Rabphilin 3A by UBE3A serves a non-degradative function"

**Authors/Affiliations**

Rossella Avagliano Trezza[1,2,5], A. Mattijs Punt[2,5], Edwin Mientjes[2], Marlene van den Berg[1], F. Isabella Zampeta[2], Ilona J. de Graaf[1,2], Yana van der Weegen[1], Jeroen A. A. Demmers[3], Ype Elgersma[2,4*], Ben Distel[1,2,4*]

[1]Department of Medical Biochemistry, Amsterdam UMC, University of Amsterdam, Amsterdam, 1105 AZ, The Netherlands

[2]Department of Neuroscience, Erasmus MC, Rotterdam, Rotterdam, 3015 CN, The Netherlands

[3]Proteomics Center, Erasmus MC, Rotterdam, 3015 CN, The Netherlands

[4]*ENCORE* Expertise Center for Neurodevelopmental Disorders, Erasmus MC, Rotterdam, Rotterdam, 3015 CN, The Netherlands

[5]These authors contributed equally

### Supplementary Fig. 1

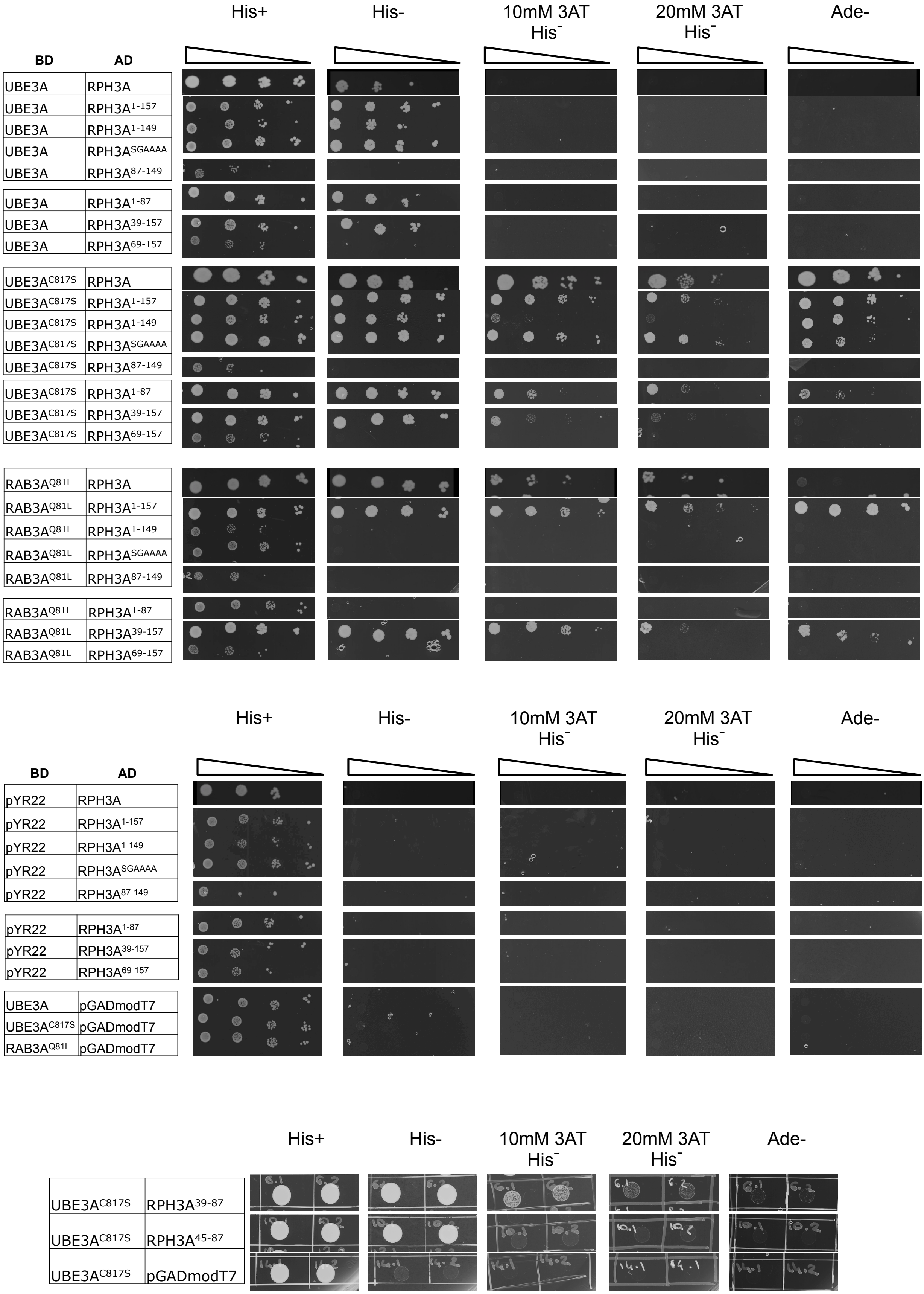


**Supplementary Figure S1**. **Identification of RPH3A as a binding partner of UBE3A.** Original plates of the spot assays corresponding to Figure 1.

### Supplementary Fig. 2

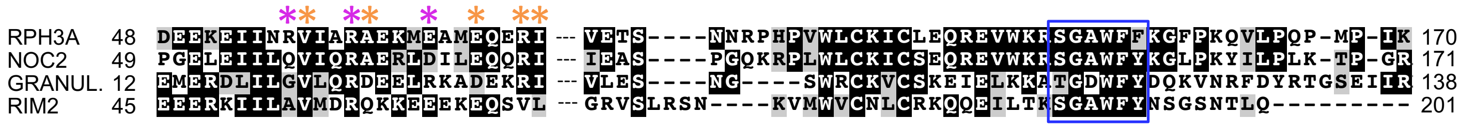


**Supplementary Figure S2. Alignment of GTPase effectors.** Multiple sequence alignment of different GTPase effectors: RPH3A, NOC2, GRANULOPHILLIN (GRANUL.) and RIM2. Two different regions of the N-terminus are shown: α-helix a1 (residues 48-71 in RPH3A) and the region C-terminal of the Zn-finger including the SGAWFF motif (residues124-170 in RPH3A). Residues that are identical in at least two proteins are shaded black, while similar residues are shaded grey. Asterisks mark residues that affect only the UBE3A interaction (magenta) or both UBE3A and RAB3A binding (orange). The blue box highlights the highly conserved SGAWFF domain.

### Supplementary Fig. 3

**A**

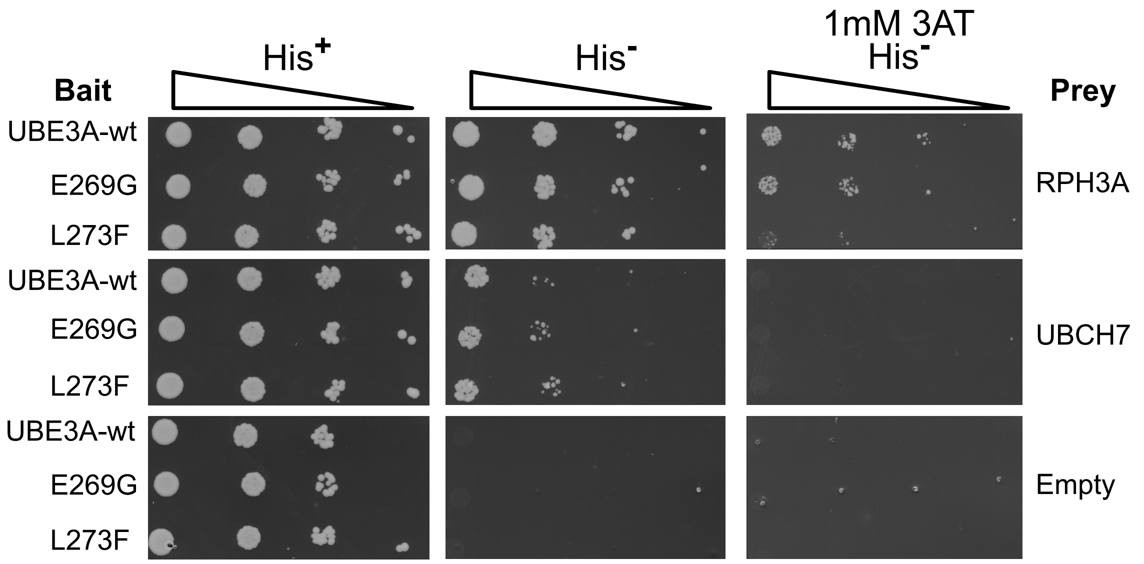


**B**

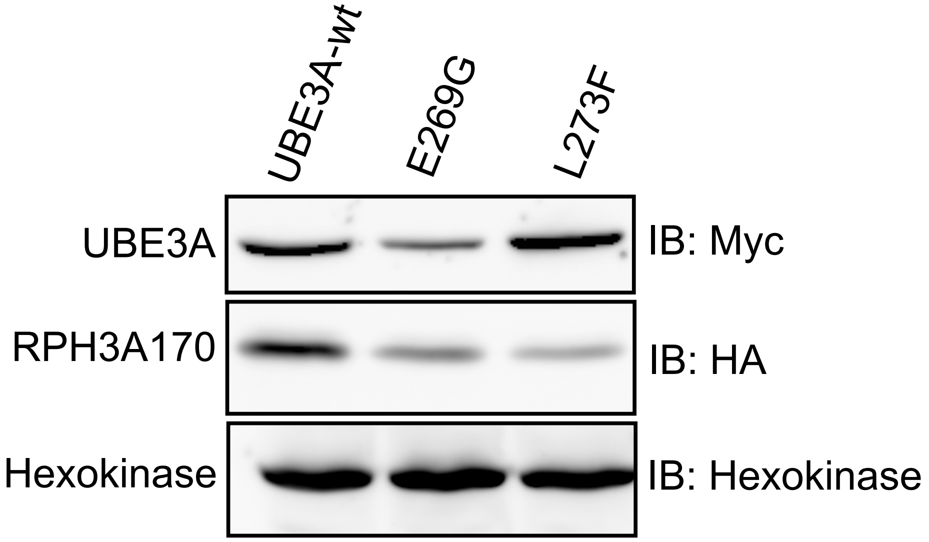


.

**Supplementary Figure S3. An AS-linked mutation in the RPH3A binding site of UBE3A interferes with RPH3A binding. a)** Original plates of the spot assay corresponding to **Figure 2c. b**) Equal expression of UBE3A variants. Total lysates obtained from yeast cells co-expressing the indicated Y2H constructs were analyzed by SDS-PAGE and immunoblotting using anti-myc (UBE3A) and anti-HA (RPH3A) antibodies. Hexokinase was used as loading control. Full-length blots are presented in **Supplementary Figure S12**

### Supplementary Fig. 4

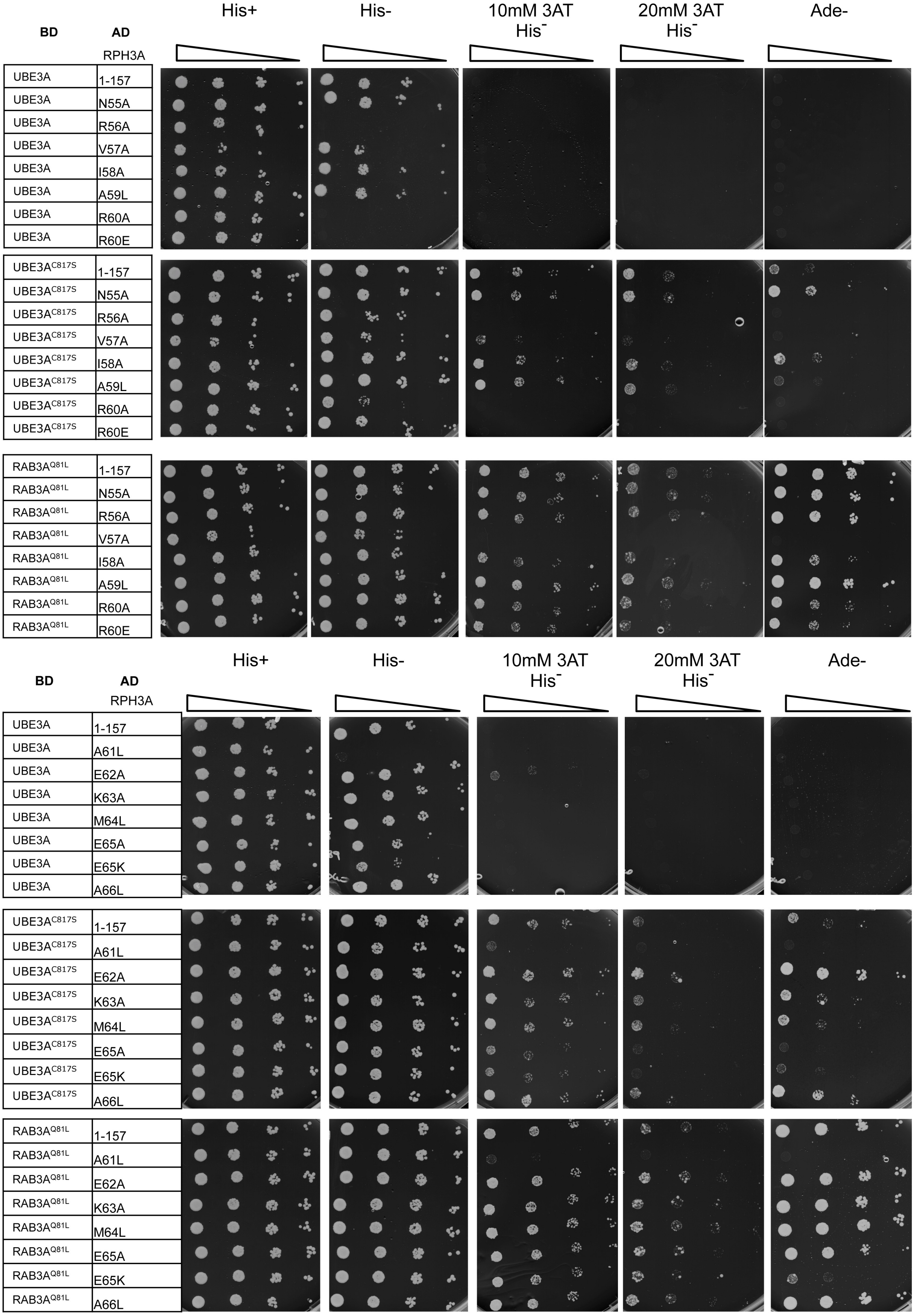


**Supplementary Figure S4. Identification of the RPH3A residues involved in UBE3A and RAB3A interaction.** Original plates of the spot assays corresponding to **Figure 3a**.

### Supplementary Fig. 4 continued

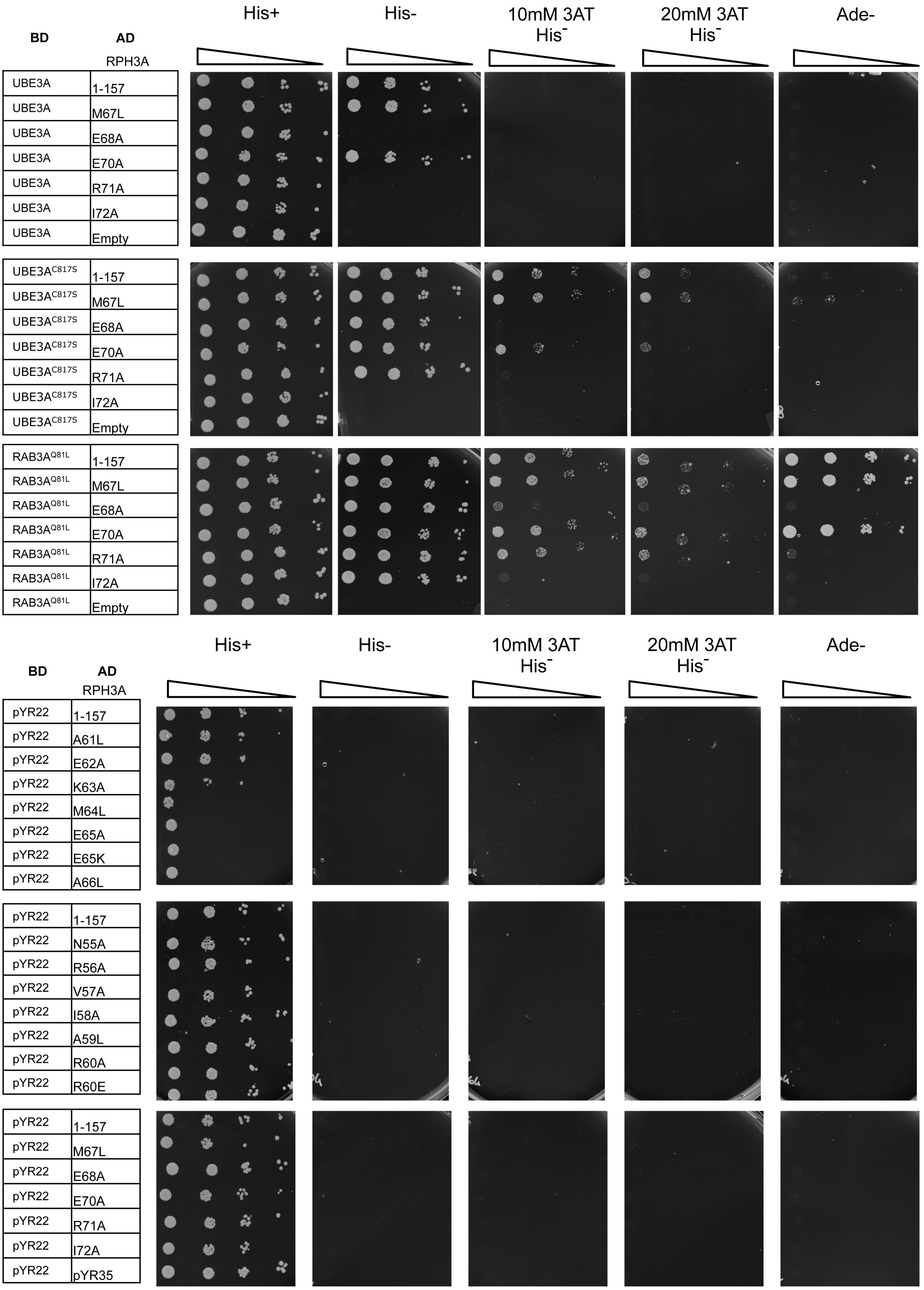


**Supplementary Figure S4. Identification of the RPH3A residues involved in UBE3A and RAB3A interaction.** Original plates of the spot assays corresponding to **Figure 3a**.

### Supplementary Figure 5

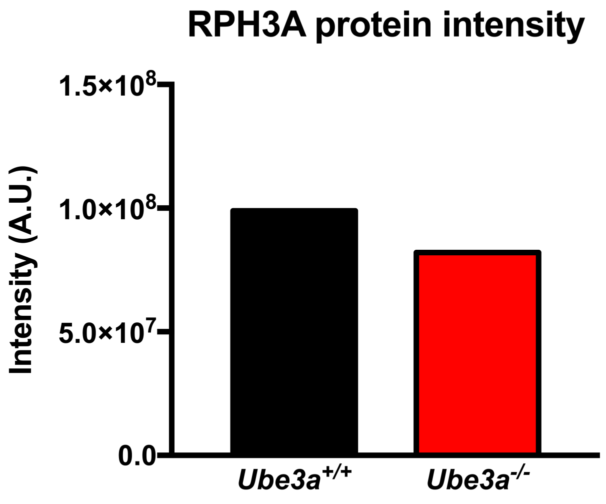


**Supplementary Figure S5. RPH3A protein levels in *Ube3a*$^{m+/p+}$ and *Ube3a*$^{m-/p-}$ mice.** RPH3A protein intensities in cortical brain lysates *Ube3a*$^{m+/p+}$ and *Ube3a*$^{m-/p-}$ mice as determined by mass spectrometry.

#### Supplementary Figure 6

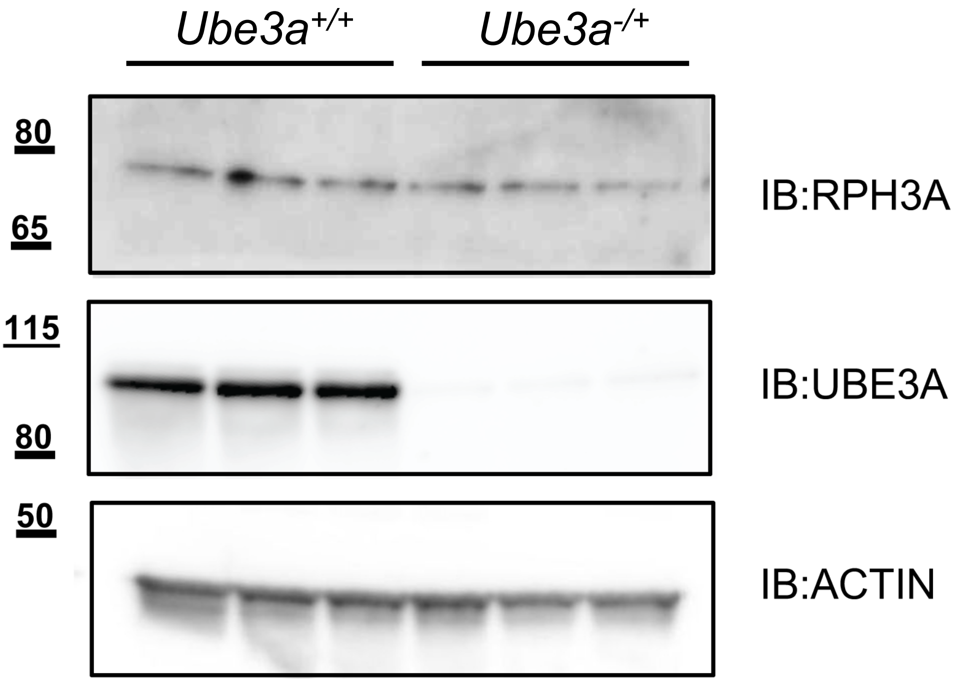


**Supplementary Figure S6. RPH3A levels are not increased in AS mice**. Cortical lysates obtained form WT (*Ube3a*$^{m+/p+}$) and AS (*Ube3a*$^{m-/p+}$) mice were analyzed by SDS-PAGE and immunoblotting using antibodies directed against RPH3A and UBE3A. ACTIN was used as loading control. Blot used for quantification as shown in **Figure 6k**.

### Supplementary Figure 7

| BD | E6 | AD |
|---|---|---|
| UBE3A | - | RPH3A170 |
| UBE3A | + | RPH3A170 |
| C817S | - | RPH3A170 |
| C817S | + | RPH3A170 |
| UBE3A | - | p53 |
| UBE3A | + | p53 |
| C817S | - | p53 |
| C817S | + | p53 |

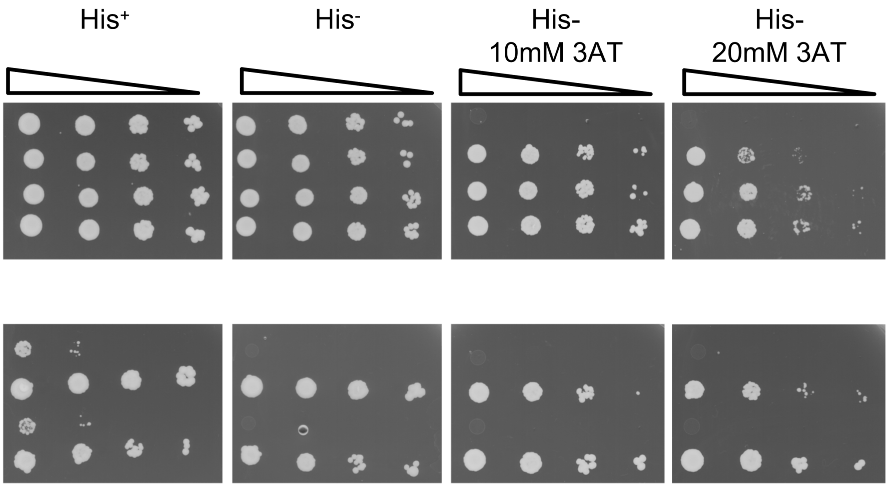


**Supplementary Figure S7. E6 enhances the RPH3A-UBE3A interaction.** *S. cerevisiae* strain PJ69a was co-transformed with UBE3A or UBE3A$^{C817S}$ (baits) and RPH3A$^{1-170}$ or p53 (preys), and non-fused E6 or empty vector (pRA1). Cells were spotted on selective plates as indicated.

### Supplementary Figure 8

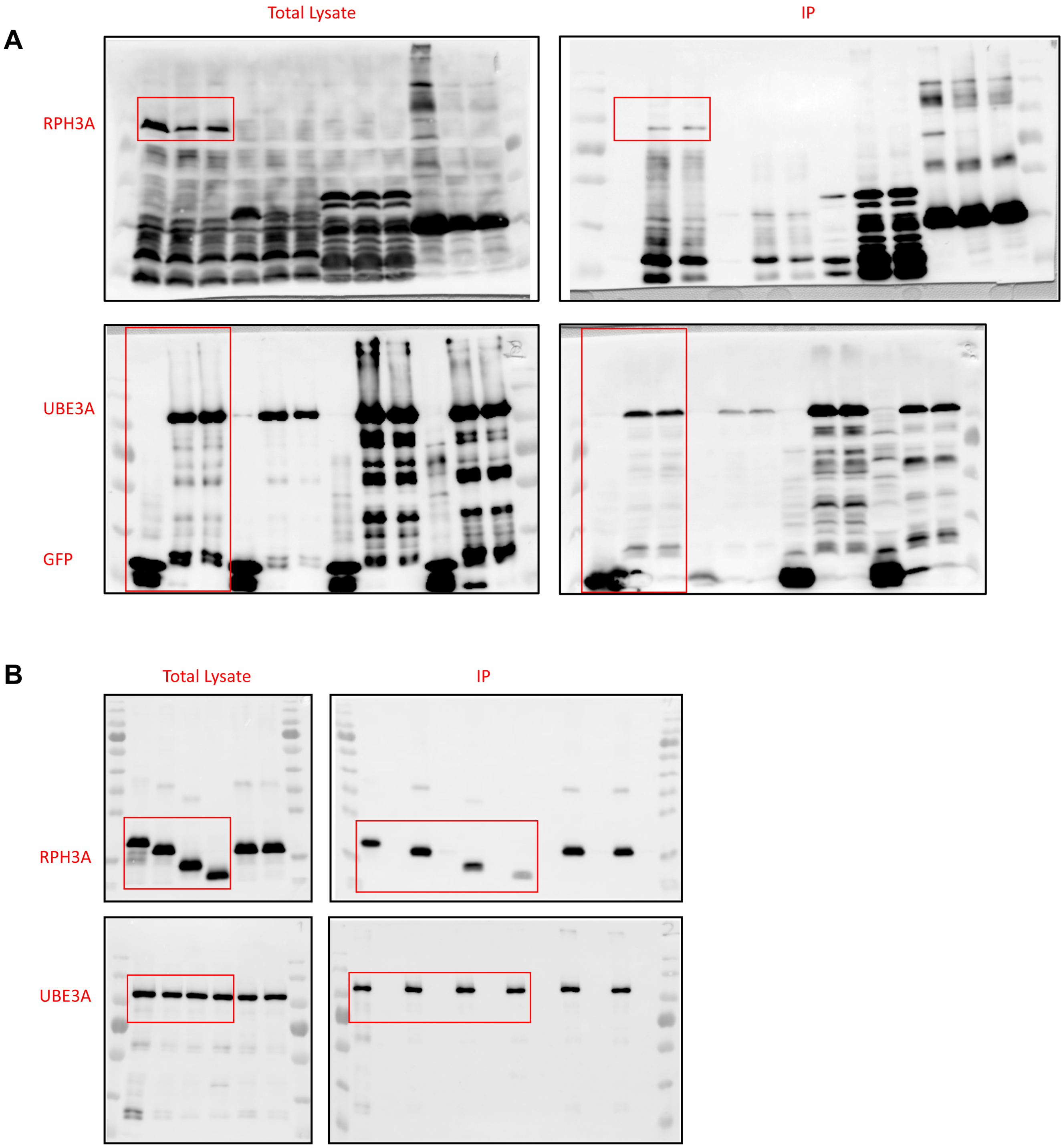


**Supplementary Figure S8. Full-length blots.** Full blots corresponding to **Figures 2a and b**

#### Supplementary Figure 9

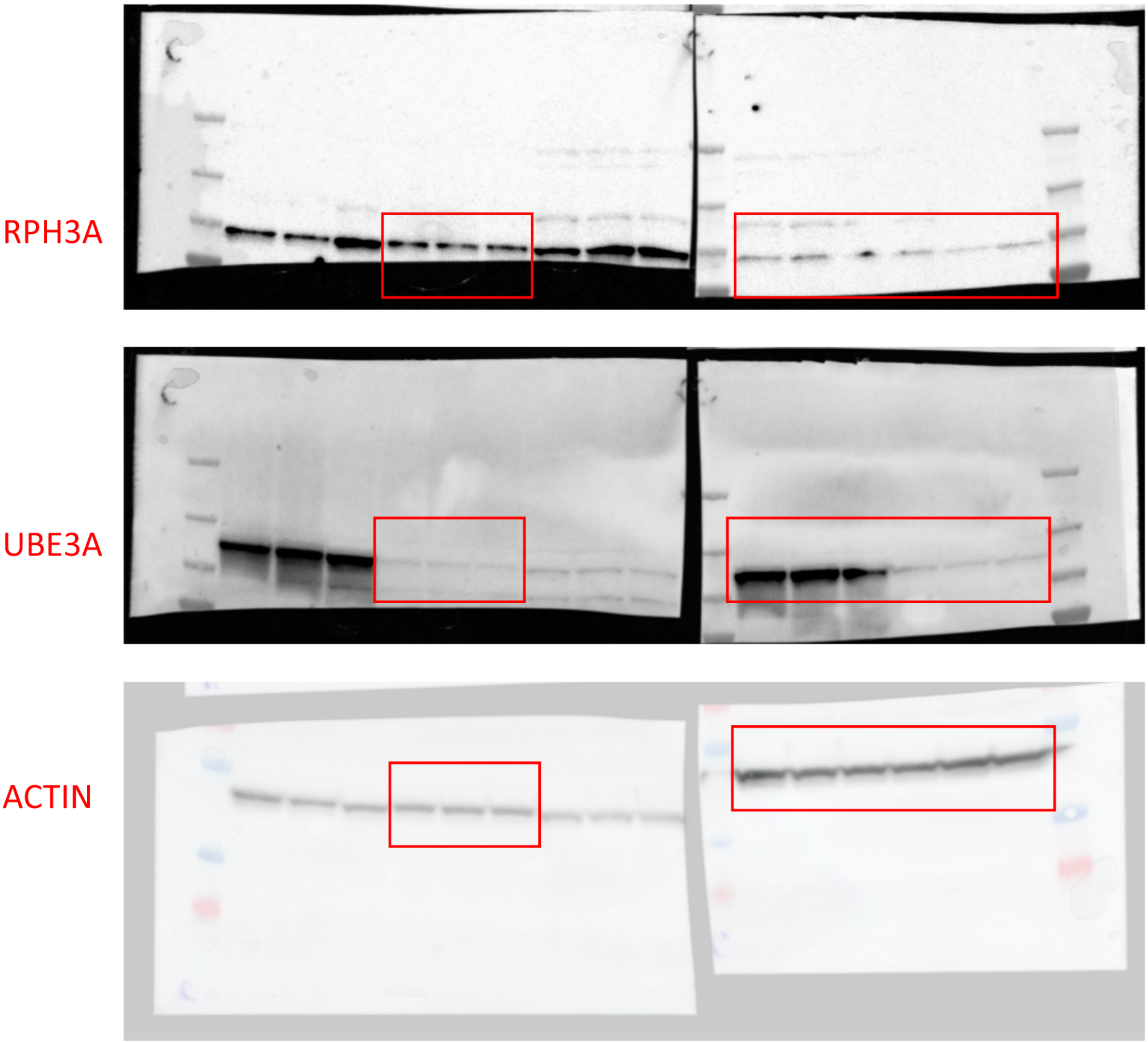


**Supplementary Figure S9. Full-length blots.** Full blots corresponding to **Figure 5a.** Blotted membranes were cut prior to antibody incubation.

### Supplementary Figure 10

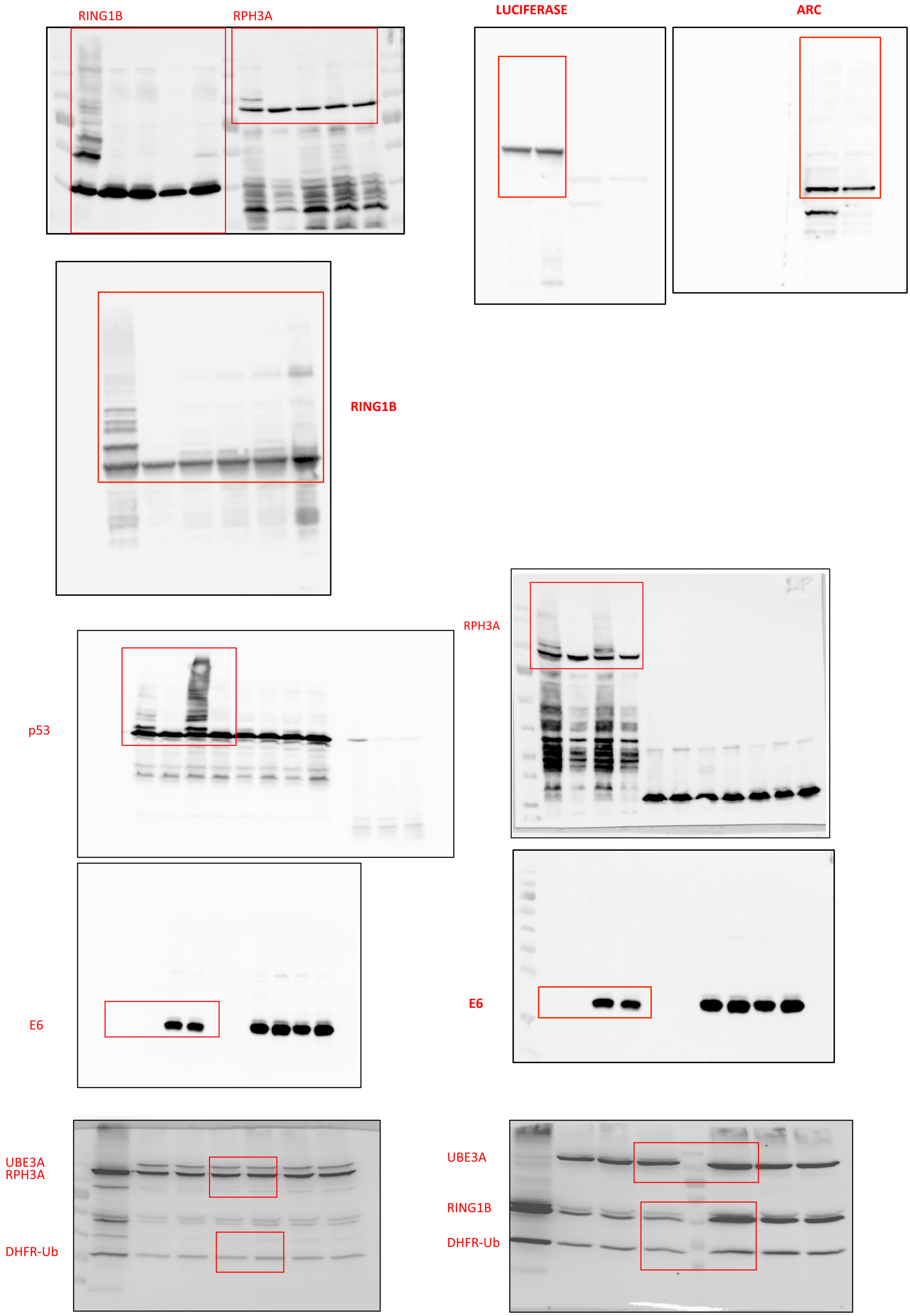


**Supplementary Figure S10. Full-length blots.** Full blots corresponding to **Figure 6d-j**

### Supplementary Figure 11

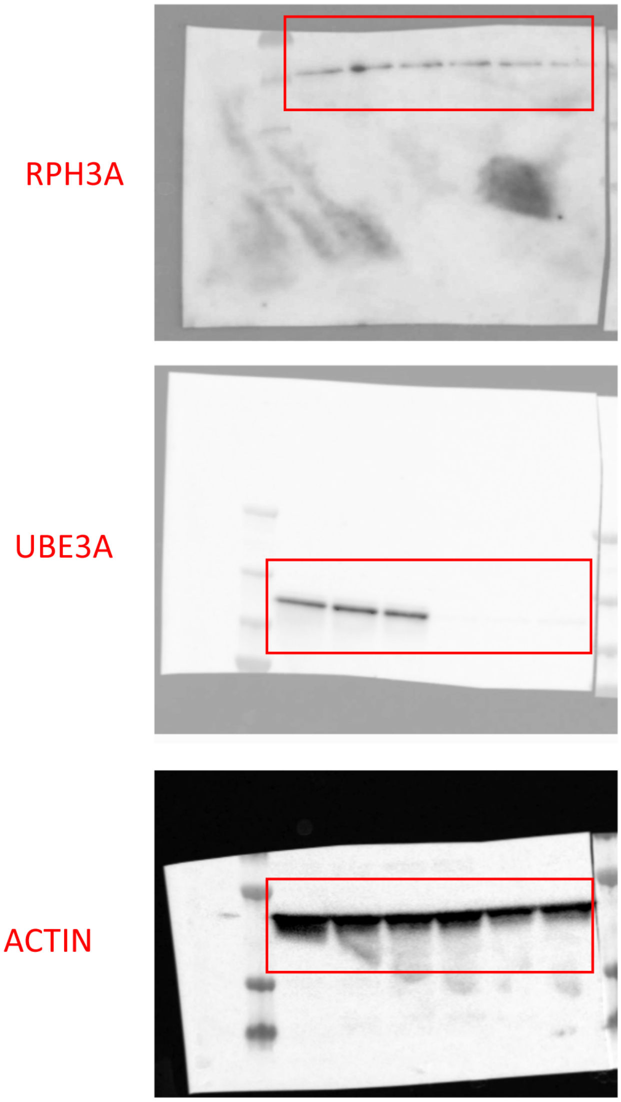


**Supplementary Figure S11. Full-length blots.** Full blots used for quantification in **Figure 6k** (see also **Supplementary Figure S6**).

### Supplementary Figure 12

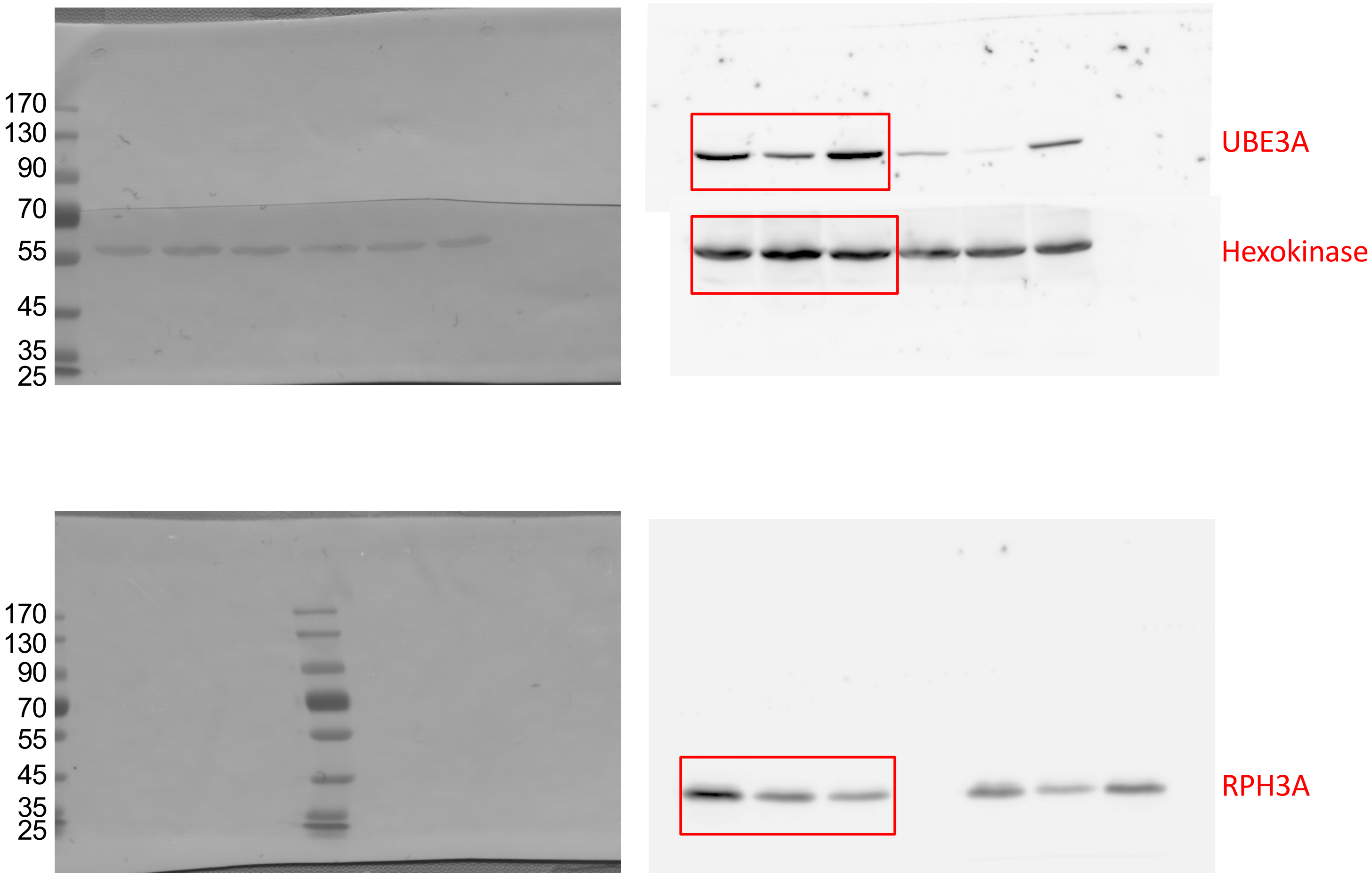


**Supplementary Figure S12. Full-length blots.** Full blots corresponding to **Figure S3b**.

**Supplementary Table 1: Primers used in this study**

| Name | Sequence | Purpose |
|---|---|---|
| p881 | GGCGCGCCTATGAAGCGAGCAGCTGCAAAGC | PCR UBE3A (Fw) |
| p882 | GTCGACTTACCTAATCACAACAGATTC | PCR UBE3A (Rv) |
| RA80 | AAGGATCCGACATGTCCGCCACAGACTCTCGCTATG | PCR RAB3A (Fw) |
| RA81 | AAGCGGCCGCGTCGACTCAGCAGGCACAATCCTGATG | PCR RAB3A (Rv) |
| RA12 | AAAGGCGCGCCTATGACTGACACTGTGGTGAAC | PCR RPH3A (Fw) |
| RA13 | AAAGTCGACCTAATCACTGGACACGTGG | PCR RPH3A FL (Rv) |
| RA101 | AAGCGGCCGCGTCGACTTAACCTTTGAAGAACCATGCTCC | PCR RPH3A 1-157 (Rv) |
| RA78 | AAGCGGCCGCGTCGACTTACTTTATAGGCATGGGCTGTGG | PCR RPH3A 1-170 (Rv) |
| RA120 | AAGGATCCGACATGTCTAGGCAGAGGAAGCAGGAAGAAC | PCR RPH3A 39-157 (Fw) |
| RA119 | AAGGATCCGACATGTCTCGCATTGGGCGCCTGGTG | PCR RPH3A 71-157 (Fw) |
| RA99 | AAGGATCCGACATGTCTGGAGATGGCGTGAACCGCT | PCR RPH3A 87-157 (Fw) |
| RA118 | AAGCGGCCGCGTCGACTTAGCGCTTCCAGACCTCTCTCT | PCR RPH3A 1-149 (Rv) |
| RA121 | AAGCGGCCGCGTCGACTTAAGCCACATTCTTCCTCATGGTC | PCR RPH3A 1-87 (Rv) |
| RA562 | AAGGATCCGACATGTCTGAACTGACAGACGAGGAG | PCR RPH3A 45-87 (Fw) |
| RA102 | CTGGAAGCGCTCAGGAGCAGCGGCAGCCAAAGGTTTCCCC AAGCAG | SDM RPH3A 1-157 SGAAAA (Fw) |
| RA103 | CTGCTTGGGGAAACCTTTGGCTGCCGCTGCTCCTGAGCGC TTCCAG | SDM RPH3A 1-157 SGAAAA (Rv) |
| RA135 | GCCAGAGTGATTGCTCGGGCAGAGAAGATGGAAGCCATGG AACAGGAACGCATTGGACGT | SDM RPH3A 1-157 N55A (Fw) |
| RA136 | CCAATGCGTTCCTGTTCCATGGCTTCCATCTTCTCTGCCCG AGCAATCACTCTGGCGATG | SDM RPH3A 1-157 N55A (Rv) |
| RA137 | AACGCAGTGATTGCTCGGGCAGAGAAGATGGAAGCCATGG AACAGGAACGCATTGGACGT | SDM RPH3A 1-157 R56A (Fw) |
| RA138 | CCAATGCGTTCCTGTTCCATGGCTTCCATCTTCTCTGCCCG AGCAATCACTGCGTTGATG | SDM RPH3A 1-157 R56A (Rv) |
| RA139 | AACAGAGCGATTGCTCGGGCAGAGAAGATGGAAGCCATGG AACAGGAACGCATTGGACGT | SDM RPH3A 1-157 V57A (Fw) |
| RA140 | CCAATGCGTTCCTGTTCCATGGCTTCCATCTTCTCTGCCCG AGCAATCGCTCTGTTGATG | SDM RPH3A 1-157 V57A (Rv) |
| RA141 | AACAGAGTGGCTGCTCGGGCAGAGAAGATGGAAGCCATGG AACAGGAACGCATTGGACGT | SDM RPH3A 1-157 I58A (Fw) |
| RA142 | CCAATGCGTTCCTGTTCCATGGCTTCCATCTTCTCTGCCCG AGCAGCCACTCTGTTGATG | SDM RPH3A 1-157 I58A (Rv) |
| RA143 | AACAGAGTGATTCTTCGGGCAGAGAAGATGGAAGCCATGGA ACAGGAACGCATTGGACGT | SDM RPH3A 1-157 A59L (Fw) |
| RA144 | CCAATGCGTTCCTGTTCCATGGCTTCCATCTTCTCTGCCCG AAGAATCACTCTGTTGATG | SDM RPH3A 1-157 A59L (Rv) |
| RA166 | AACAGAGTGATTGCTGCTGCAGAGAAGATGGAAGCCATGGA ACAGGAACGCATTGGACGT | SDM RPH3A 1-157 R60A (Fw) |
| RA167 | CCAATGCGTTCCTGTTCCATGGCTTCCATCTTCTCTGCAGCA GCAATCACTCTGTTGATG | SDM RPH3A 1-157 R60A (Rv) |
| RA147 | AACAGAGTGATTGCTGAGGCAGAGAAGATGGAAGCCATGG AACAGGAACGCATTGGACGT | SDM RPH3A 1-157 R60E (Fw) |
| RA148 | CCAATGCGTTCCTGTTCCATGGCTTCCATCTTCTCTGCCTCA GCAATCACTCTGTTGATG | SDM RPH3A 1-157 R60E (Rv) |
| RA168 | AACAGAGTGATTGCTCGGTTGGAGAAGATGGAAGCCATGGA ACAGGAACGCATTGGACGT | SDM RPH3A 1-157 A61L (Fw) |

| | | |
|---|---|---|
| RA170 | CCAATGCGTTCCTGTTCCATGGCTTCCATCTTCTCCAACCGAGCAATCACTCTGTTGATG | SDM RPH3A 1-157 A61L (Rv) |
| RA171 | AACAGAGTGATTGCTCGGGCAGCTAAGATGGAAGCCATGGAACAGGAACGCATTGGACGT | SDM RPH3A 1-157 E62A (Fw) |
| RA172 | CCAATGCGTTCCTGTTCCATGGCTTCCATCTTAGCTGCCCGAGCAATCACTCTGTTGATG | SDM RPH3A 1-157 E62A (Rv) |
| RA173 | AACAGAGTGATTGCTCGGGCAGAGGCTATGGAAGCCATGGAACAGGAACGCATTGGACGT | SDM RPH3A 1-157 K63A (Fw) |
| RA174 | CCAATGCGTTCCTGTTCCATGGCTTCCATAGCCTCTGCCCGAGCAATCACTCTGTTGATG | SDM RPH3A 1-157 K63A (Rv) |
| RA175 | AACAGAGTGATTGCTCGGGCAGAGAAGTTGGAAGCCATGGAACAGGAACGCATTGGACGT | SDM RPH3A 1-157 M64A (Fw) |
| RA176 | CCAATGCGTTCCTGTTCCATGGCTTCCAACTTCTCTGCCCGAGCAATCACTCTGTTGATG | SDM RPH3A 1-157 M64A (Rv) |
| RA149 | AACAGAGTGATTGCTCGGGCAGAGAAGATGAAAGCCATGGAACAGGAACGCATTGGACGT | SDM RPH3A 1-157 E65K (Fw) |
| RA150 | CCAATGCGTTCCTGTTCCATGGCTTTCATCTTCTCTGCCCGAGCAATCACTCTGTTGATG | SDM RPH3A 1-157 E65K (Rv) |
| RA177 | AACAGAGTGATTGCTCGGGCAGAGAAGATGGCTGCCATGGAACAGGAACGCATTGGACGT | SDM RPH3A 1-157 E65A (Fw) |
| RA178 | CCAATGCGTTCCTGTTCCATGGCAGCCATCTTCTCTGCCCGAGCAATCACTCTGTTGATG | SDM RPH3A 1-157 E65A (Rv) |
| RA179 | AACAGAGTGATTGCTCGGGCAGAGAAGATGGAATTGATGGAACAGGAACGCATTGGACGT | SDM RPH3A 1-157 A66L (Fw) |
| RA180 | CCAATGCGTTCCTGTTCCATCAATTCCATCTTCTCTGCCCGAGCAATCACTCTGTTGATG | SDM RPH3A 1-157 A66L (Rv) |
| RA181 | AACAGAGTGATTGCTCGGGCAGAGAAGATGGAAGCCTTGGAACAGGAACGCATTGGACGT | SDM RPH3A 1-157 M67L (Fw) |
| RA182 | CCAATGCGTTCCTGTTCCAAGGCTTCCATCTTCTCTGCCCGAGCAATCACTCTGTTGATG | SDM RPH3A 1-157 M67L (Rv) |
| RA183 | AACAGAGTGATTGCTCGGGCAGAGAAGATGGAAGCCATGGCTCAGGAACGCATTGGACGT | SDM RPH3A 1-157 E68A (Fw) |
| RA184 | CCAATGCGTTCCTGAGCCATGGCTTCCATCTTCTCTGCCCGAGCAATCACTCTGTTGATG | SDM RPH3A 1-157 E68A (Rv) |
| RA187 | AACAGAGTGATTGCTCGGGCAGAGAAGATGGAAGCCATGGAACAGGCTCGCATTGGACGT | SDM RPH3A 1-157 E70A (Fw) |
| RA188 | CCAATGCGAGCCTGTTCCATGGCTTCCATCTTCTCTGCCCGAGCAATCACTCTGTTGATG | SDM RPH3A 1-157 E70A (Rv) |
| RA189 | AACAGAGTGATTGCTCGGGCAGAGAAGATGGAAGCCATGGAACAGGAAGCTATTGGACGT | SDM RPH3A 1-157 R71A (Fw) |
| RA190 | CCAATAGCTTCCTGTTCCATGGCTTCCATCTTCTCTGCCCGAGCAATCACTCTGTTGATG | SDM RPH3A 1-157 R71A (Rv) |
| RA145 | AACAGAGTGATTGCTCGGGCAGAGAAGATGGAAGCCATGGAACAGGAACGCGCTGGACGT | SDM RPH3A 1-157 I72A (Fw) |
| RA146 | CCAGCGCGTTCCTGTTCCATGGCTTCCATCTTCTCTGCCCGAGCAATCACTCTGTTGATG | SDM RPH3A 1-157 I72A (Rv) |
| RA82 | GGGACACAGCAGGGCTCGAGCGGTACCGCACC | SDM RAB3A Q81L (Fw) |
| RA83 | GGTGCGGTACCGCTCGAGCCCTGCTGTGTCCC | SDM RAB3A Q81L (Rv) |
| MB3000 | GGCCGCaaggagatatacaATGGCTGAACAAAAATTGATTTCTGAAGAGGATTTGTCTGGTGCTTCTgacgtcCTGCAGGctGTTTAAACttaattaaC | construction rbs-Myc-MCS (dsOligo Fw) |
| MB3001 | ctagGttaattaaGTTTAAACagCCTGCAGgacgtcAGAAGCACCAGACAAATCCTCTTCAGAAATCAATTTTTGTTCAGCCATtgtatatctccttGC | construction rbs-Myc-MCS (dsOligo Rv) |

| | | |
|---|---|---|
| MB3002 | CCTGCAGGctATGtttcaggacccacaggagcg | PCR E6 type 16 (Fw) |
| MB3003 | ttaattaaTTACAGCTGGGTTTCTCTACGTG | PCR E6 type 16 (Rv) |
| MB3004 | AATTCgtTAAgAGGCCTgTAAgcTAAgg | deletion E2 (Fw) |
| MB3005 | CGCGccTTAgcTTAcAGGCCTcTTAacG | deletion E2 Rv) |
| MB3006 | CGCGCCtTAAggTAAgcTAAgacgt | deletion E1 (Fw) |
| MB3007 | cTTAgcTTAccTTAaGG | deletion E1 (Rv) |
| MB3008 | CTAGAAATAATTTTGTTTAACTTTAAG | deletion Ub (Fw) |
| MB3009 | aattCTTAAAGTTAAACAAAATTATTT | deletion Ub (Rv) |
| p5098 | GGCGCGCCGATGTGCAATACCAACATGTCTGTACCTAC | cloning MDM2 (Fw) |
| p5099 | AATTAAGTGCGGCCGCTCGACCCG | cloning MDM2 (Rv) |
| p3991 | GGCGCGCCTGTGGAAGACGCCAAAAACATAAAGAAA | cloning Luciferase (Fw) |
| p3992 | GCGGCCGCTTACACGGCGATCTTTC | cloning Luciferase (Rv) |
| p5131 | CGCGCCAACGTCTGGCGCGGGTAC | cloning NEDD4 (Fw) |
| p5132 | CCGCGCCAGACGTTGG | cloning NEDD4 (Rv) |
| RA220 | CTCAATTTGTTCATTATTGTAATGGGGAATAGTAATCTCCAC AGTCCTGAATATC | SDM E269G (Fw) |
| RA221 | GATATTCAGGACTGTGGAGATTACTATTCCCCATTACAAT AATGAACAAATTGAG | SDM E269G (Rv) |
| RA224 | CATTATTGTAATGGAGAATAGTAATTTCCACAGTCCTGAA TATCTGGAAATG | SDM L273F (Fw) |
| RA225 | CATTTCCAGATATTCAGGACTGTGGAAATTACTATTCTC CAT TACAATAATG | SDM L273F (Rv) |

**Supplementary Table 2: Plasmids generated and used in this study**

| Name | Insert | Purpose |
|---|---|---|
| pYR22 | Empty | Y2H |
| pGADT7 | Empty | Y2H |
| pYR35 | Empty | Y2H |
| pRA55 | UBE3A WT | Y2H |
| pRA61 | UBE3A C817S | Y2H |
| pRA72 | RAB3A Q81L | Y2H |
| pRA28 | RPH3A FL | Y2H |
| pRA77 | RPH3A 1-157 | Y2H |
| pRA87 | RPH3A 1-157 SGAAAA | Y2H |
| pRA98 | RPH3A 39-157 | Y2H |
| pRA99 | RPH3A 71-157 | Y2H |
| pRA89 | RPH3A 87-149 | Y2H |
| pRA86 | RPH3A 1-149 | Y2H |
| pRA100 | RPH3A 1-87 | Y2H |
| pRA106 | RPH3A 39-87 | Y2H |
| pRA110 | RPH3A 45-87 | Y2H |
| pMB277 | E1, UbcH7, Ub wt | *E. coli* expression |
| pMB278 | E1, UbcH7, ΔUb | *E. coli* expression |
| pMB280 | V5-PEX22, HA-GFP | *E. coli* expression |
| pMB284 | V5-RPH3A FL, HA-GFP | *E. coli* expression |
| pMB286 | V5-RPH3A FL, HA-UBE3A WT | *E. coli* expression |
| pMB287 | V5-RPH3A FL, HA-UBE3A C817S | *E. coli* expression |
| pMB316 | E1, ΔE2, Ub | *E. coli* expression |
| pMB318 | ΔE1, UbcH7, Ub | *E. coli* expression |
| pMB323 | V5-PEX22, HA-GFP, Myc/MCS | *E. coli* expression |
| pMB326 | V5-RPH3A 1-157, HA-UBE3A WT | *E. coli* expression |
| pMB327 | V5-RPH3A 39-157, HA-UBE3A WT | *E. coli* expression |
| pMB328 | V5-RPH3A 71-157, HA-UBE3A WT | *E. coli* expression |
| pMB336 | V5-PEX22, HA-GFP, Myc-E6 | *E. coli* expression |
| pMB338 | V5-RPH3A FL, HA-UBE3A WT, Myc-E6 | *E. coli* expression |
| pMB344 | V5-p53, HA-UBE3A WT | *E. coli* expression |
| pMB345 | V5-p53, HA-UBE3A C817S | *E. coli* expression |
| pMB346 | V5-p53, HA-UBE3A WT, Myc-E6 | *E. coli* expression |
| pMB347 | V5-p53, HA-UBE3A C817S, Myc-E6 | *E. coli* expression |
| pMB419 | V5/MCS | *E. coli* expression |
| pMB427 | V5-RPH3A FL | *E. coli* expression |
| pYW5 | HA/MCS-Myc/MCS | *E. coli* expression |
| pMB436 | HA-UBE3A WT | *E. coli* expression |
| pMB437 | HA-UBE3A C817S | *E. coli* expression |
| pMB443 | V5-RING1B I53S | *E. coli* expression |
| pRA172 | RPH3A 1-157 N55A | Y2H |
| pRA173 | RPH3A 1-157 R56A | Y2H |

| | | |
|---|---|---|
| pRA174 | RPH3A 1-157 V57A | Y2H |
| pRA175 | RPH3A 1-157 I58A | Y2H |
| pRA176 | RPH3A 1-157 A59L | Y2H |
| pRA141 | RPH3A 1-157 R60A | Y2H |
| pRA177 | RPH3A 1-157 R60E | Y2H |
| pRA131 | RPH3A 1-157 A61L | Y2H |
| pRA142 | RPH3A 1-157 E62A | Y2H |
| pRA143 | RPH3A 1-157 K63A | Y2H |
| pRA132 | RPH3A 1-157 M64A | Y2H |
| pRA178 | RPH3A 1-157 E65K | Y2H |
| pRA133 | RPH3A 1-157 E65A | Y2H |
| pRA134 | RPH3A 1-157 A66L | Y2H |
| pRA135 | RPH3A 1-157 M67L | Y2H |
| pRA136 | RPH3A 1-157 E68A | Y2H |
| pRA137 | RPH3A 1-157 E70A | Y2H |
| pRA138 | RPH3A 1-157 R71A | Y2H |
| pRA179 | RPH3A 1-157 I72A | Y2H |
| pRA158 | UBE3A E269G | Y2H |
| pRA159 | UBE3A L273F | Y2H |
| pRA1 | Empty | Y3H |
| pRA117 | RAB3A Q81L | Y3H |
| pRA125 | RPH3a 1-170 | Y3H |
| pRA126 | RPH3A 1-157 SGAAAA | Y3H |
| pRA53 | UBE3A WT | *S. cerevisiae* expression |
| pRA62 | UBE3A C817S | *S. cerevisiae* expression |
| pRA34 | RPH3A FL | *S. cerevisiae* expression |
| pRA64 | UBE3A WT | HEK293T expression |
| pRA65 | UBE3A C817S | HEK293T expression |
| pRA66 | RPH3A FL | HEK293T expression |
| pMS1 | RING1B I53S | HEK293T expression |
| pYE2009 | V5-ARC | *E. coli* expression |
| pYE2010 | V5-Luciferase (firefly) | *E. coli* expression |
| pYE2011 | HA-MDM2 (human) | *E. coli* expression |
| pYE2012 | HA-NEDD4 (human) | *E. coli* expression |
